# An interchangeable prion-like domain is required for Ty1 retrotransposition

**DOI:** 10.1101/2023.02.27.530227

**Authors:** Sean L. Beckwith, Emily J. Nomberg, Abigail C. Newman, Jeannette V. Taylor, Ricardo C. Guerrero, David J. Garfinkel

**Author notes:** Author contributions: S.L.B. and D.J.G. designed research, analyzed data, and wrote the paper; S.L.B., D.J.G., E.J.N., A.C.N, and J.V.T. performed research; J.V.T. and R.C.G. conducted transmission electron microscopy; S.L.B., R.C.G., and D.J.G. acquired funding.

## Abstract

Retrotransposons and retroviruses shape genome evolution and can negatively impact genome function. *Saccharomyces cerevisiae* and its close relatives harbor several families of LTR-retrotransposons, the most abundant being Ty1 in several laboratory strains. The cytosolic foci that nucleate Ty1 virus-like particle (VLP) assembly are not well-understood. These foci, termed retrosomes or T-bodies, contain Ty1 Gag and likely Gag-Pol and the Ty1 mRNA destined for reverse transcription. Here, we report a novel intrinsically disordered N-terminal <u>pr</u>ion-like <u>d</u>omain (PrLD) within Gag that is required for transposition. This domain contains amino-acid composition similar to known yeast prions and is sufficient to nucleate prionogenesis in an established cell-based prion reporter system. Deleting the Ty1 PrLD results in dramatic VLP assembly and retrotransposition defects but does not affect Gag protein level. Ty1 Gag chimeras in which the PrLD is replaced with other sequences, including yeast and mammalian prionogenic domains, display a range of retrotransposition phenotypes from wildtype to null. We examine these chimeras throughout the Ty1 replication cycle and find that some support retrosome formation, VLP assembly, and retrotransposition, including the yeast Sup35 prion and the mouse PrP prion. Our interchangeable Ty1 system provides a useful, genetically tractable *in vivo* platform for studying PrLDs, complete with a suite of robust and sensitive assays, and host modulators developed to study Ty1 retromobility. Our work invites study into the prevalence of PrLDs in additional mobile elements.

**Significance:** Retrovirus-like retrotransposons help shape the genome evolution of their hosts and replicate within cytoplasmic particles. How their building blocks associate and assemble within the cell is poorly understood. Here, we report a novel <u>pr</u>ion-like <u>d</u>omain (PrLD) in the budding yeast retrotransposon Ty1 Gag protein that builds virus-like particles. The PrLD has similar sequence properties to prions and disordered protein domains that can drive the formation of assemblies that range from liquid to solid. We demonstrate that the Ty1 PrLD can function as a prion and that certain prion sequences can replace the PrLD and support Ty1 transposition. This interchangeable system is an effective platform to study additional disordered sequences in living cells.

## Introduction

Retrotransposons are pervasive across diverse eukaryotes and influence genome evolution and affect host fitness. The budding yeast *Saccharomyces cerevisiae* contains Ty1-5 long-terminal repeat (LTR)-retrotransposons, with Ty1 as the most abundant element in many laboratory strains (1, 2). LTR-retrotransposons are the evolutionary progenitors of retroviruses; Ty1 elements share many structural hallmarks with retroviral genomic RNA and undergo an analogous replication cycle but lack an extracellular phase. Ty1 is transcribed from LTR-to-LTR and contains two partially overlapping open reading frames: *GAG* and *POL*. Ty1 RNA serves as a template for protein synthesis and reverse transcription. Translation of Ty1 *POL* requires a programmed +1 frameshift near the C-terminus of *GAG*, resulting in a large Gag-Pol precursor (p199) (3). Like retroviral RNA, Ty1 RNA is specifically packaged into virus-like particles (VLPs) where RNA is present in a dimeric form (4–7). Proteolytic protein maturation occurs within VLPs by a protease (PR) encoded within *GAG* and *POL*. Ty1 PR cleaves the Gag-p49 precursor near the C-terminus to generate p45, the capsid protein, and Gag-Pol-p199 to form mature PR, integrase (IN), and reverse transcriptase (RT) (8, 9). Reverse transcription occurs within mature VLPs and, like HIV-1, requires a complex formed between RT and IN (10, 11). Ty1 preferentially integrates upstream of genes actively transcribed by RNA Polymerase III (Pol III) due to interactions between IN and Pol III subunits (12–14).

Ty1 Gag performs the same functions as retroviral capsid and nucleocapsid. Amino acids 159-355 encode NTD and CTD capsid folds, assembling VLPs (15), and a C-terminal domain of Gag displays nucleic acid chaperone (NAC) activity (16, 17). Sequences in the Ty1 RNA encoding the Gag protein are required for packing, dimerization, and reverse transcription (3). The N-terminal region of Gag has unknown function, and it is not known whether it is required for transposition.

While several steps of retrotransposon life cycles have been investigated, it is not well-understood how their RNA genomes and protein machinery associate within the cellular milieu to facilitate VLP assembly and replication. Retroviral particle assembly often occurs in subcellular domains, referred to as “viral factories” or “viral inclusions” (18, 19). The sites of Ty1 VLP assembly are cytoplasmic foci termed retrosomes, or T-bodies, which contain Ty1 RNA, Gag, Gag-Pol, and perhaps additional cellular proteins (20–22). What drives the biogenesis of retrosomes is not understood. Mounting evidence suggests liquid-liquid phase separation (LLPS) underlies many examples of membraneless compartments (23, 24). Aggregation-prone proteins that drive LLPS have overlapping properties with prions, and both are implicated in age-related disease (25–34). Spontaneous demixing in these systems is often facilitated by intrinsically disordered domains, multivalent proteins, and scaffolding around nucleic acids. Indeed, prion-like and LLPS mechanisms provide intriguing models for retroelement assembly steps. Ty1 retrosomes contain Ty1 RNA and Gag oligomers associated with the RNA. Several viruses utilize LLPS in replication and assembly, including rabies virus (35), influenza A (36), herpes simplex virus 1 (37), measles virus (38), HIV-1 (39), and SARS-CoV-2 (40). Also, the human retrotransposon LINE-1 has been reported to phase separate *in vitro* (41). Here, we present evidence that the Ty1 Gag protein contains a prion-like domain required for VLP assembly and transposition, raising the possibility that Ty1 Gag facilitates prion-like or phase separating behaviors within retrosomes.

## Results

### Bioinformatic analyses reveal a prion-like domain in Ty1 Gag

Ty1 Gag contains several protein features, including capsid and nucleic acid chaperone domains (15, 17). The N-terminal region of the protein, meanwhile, is predicted to be unstructured and does not have previously reported function. We analyzed Ty1 Gag (Fig. 1*A*) and Gag-Pol (Fig. S1) using several bioinformatic tools designed to predict protein disorder, amyloidogenic secondary structures, and amino acid composition similarity to known yeast prions (42–45). For comparison, we ran the well-studied yeast prions Sup35 and Ure2, the mouse prion protein PrP, and Alzheimer’s disease-associated human Aβ_1-42_ through the same bioinformatic analyses (Fig. 1*B-E*). Ty1 Gag contains a 71-amino acid domain with strikingly similar amino acid composition to yeast prions in its disordered N-terminus, comparable to Sup35 and Ure2. This Gag <u>pr</u>ion-like <u>d</u>omain (PrLD) is predicted to be unstructured by AlphaFold (46) and no published structures of the region are available, similar to canonical prions (15, 47–50) (Fig. S2). Given the computational predictions and the requirement for Gag in forming Ty1 retrosomes, we further investigated prionogenic properties of the Gag PrLD, which we define as amino acid residues 66-136.

**Fig 1.**
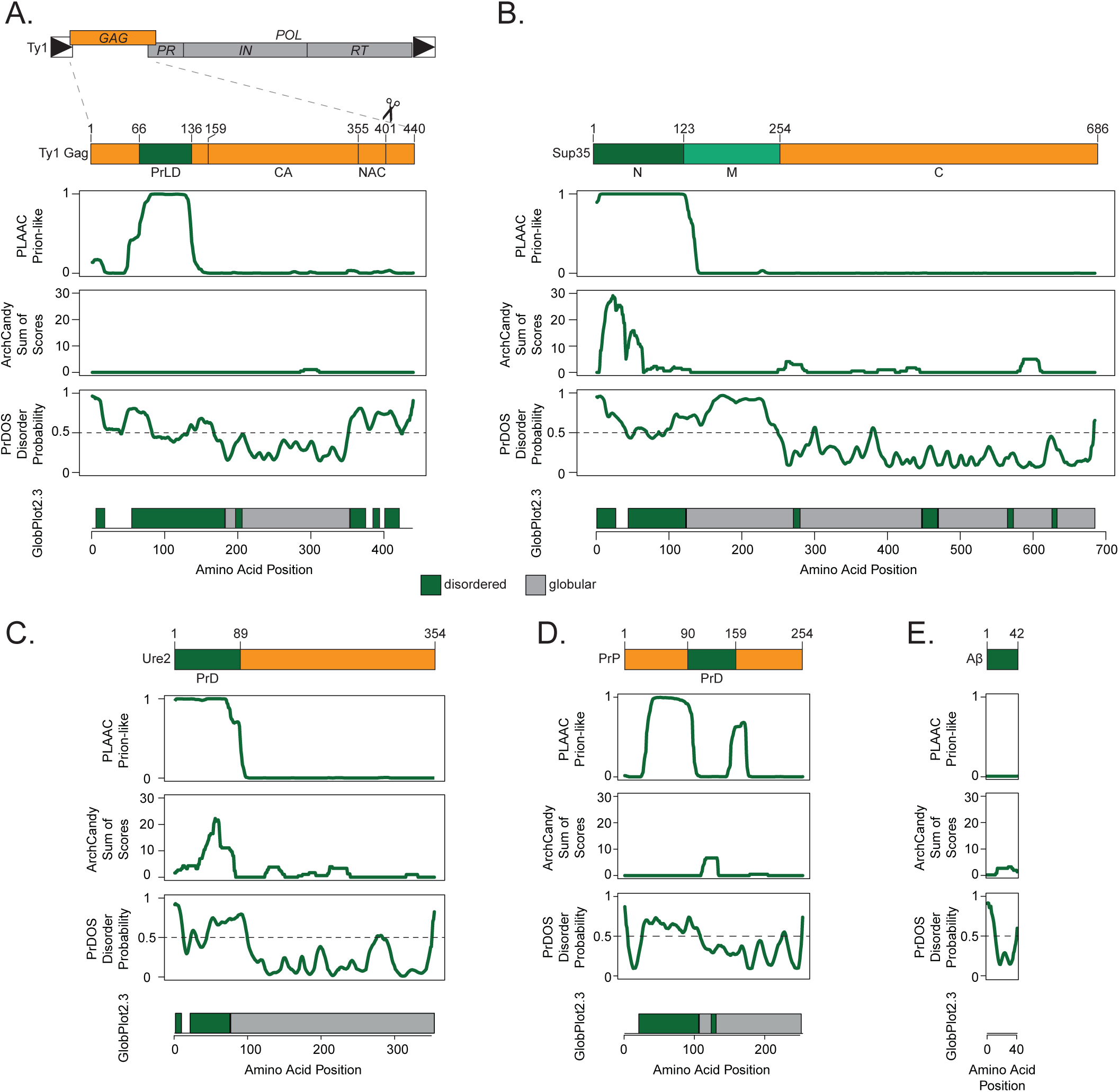
The Ty1 retrotransposon Gag contains a prion-like domain. Schematic of the Ty1 retrotransposon gene organization, with a detailed view of domains of the Gag protein (*A*), yeast prion Sup35 (*B*), yeast prion Ure2 (*C*), mouse prion protein (PrP) (*D*), and human amyloid beta (Aβ) (*E*); PrLD = prion-like domain, capsid domain (CA) and nucleic acid chaperone domain (NAC) are defined in (15). Below are bioinformatic analyses of each protein aligned with the schematic above: yeast prion-like amino acid composition (PLAAC), predicted amyloidogenic regions (ArchCandy), predicted protein disorder (PrDOS), predicted disordered (green) and globular (grey) regions (GlobPlot2.3).

### Prionogenic properties of the Gag_PrLD_

We used a well-characterized Sup35-based *in vivo* reporter system to assess the ability of the Gag PrLD to promote prionogenesis in a yeast strain harboring a mutant allele of the adenine biosynthesis gene, *ade1-14*, which contains a premature stop codon (Fig. 2*A*) (51–53). Soluble Sup35 functions as a translation termination factor, resulting in a truncated non-functional Ade1 protein. Yeast fails to grow on media lacking adenine and appears red due to the buildup of a metabolic intermediate. However, formation of a prion state (termed [*PSI*^+^]) aggregates Sup35 away from the ribosome, allowing for translational readthrough. This can be detected by adenine prototrophy and yeast colonies appearing white. Fusion of the PrLD of interest to the Sup35 N or NM domains promotes prion nucleation and has previously been used to study mammalian PrP and Aβ (53). Expression of Sup35NM-Gag_PrLD_ fusion under the *CUP1* copper-inducible promoter stimulates prionogenesis, as detected by increased papillation on SC-Ade when compared to the reporter alone (Fig. 2*B*). Growth is copper responsive, however, we found that the Sup35N reporter construct displays a high background growth independent of induction (Fig. S3*A-B*). We next biochemically monitored prion aggregation using semi-denaturing detergent-agarose gel electrophoresis (SDD-AGE) (54). Gag_PrLD_ fusions formed large, slow-migrating, copper-inducible aggregates with both Sup35N (Fig. 2*C*) and Sup35NM (Fig. S2*C*) above reporter alone. Finally, we verified prion nucleation specifically, as opposed to colony growth due to accumulating suppressor mutations, by curing colonies of the prion after passaging cells on guanidine hydrochloride (GdHCl) (55, 56). Representative cells are shown for the naïve [*psi*^-^], induced [*PSI*^+^], and cured states for Sup35NM fusions to Gag_PrLD_ or positive control Aβ, both HA-tagged (Fig. 2*D*) and untagged (Fig. S2*D*). A large fraction of Sup35NM-Gag_PrLD_ Ade^+^ colonies were curable by GdHCl.

**Fig 2.**
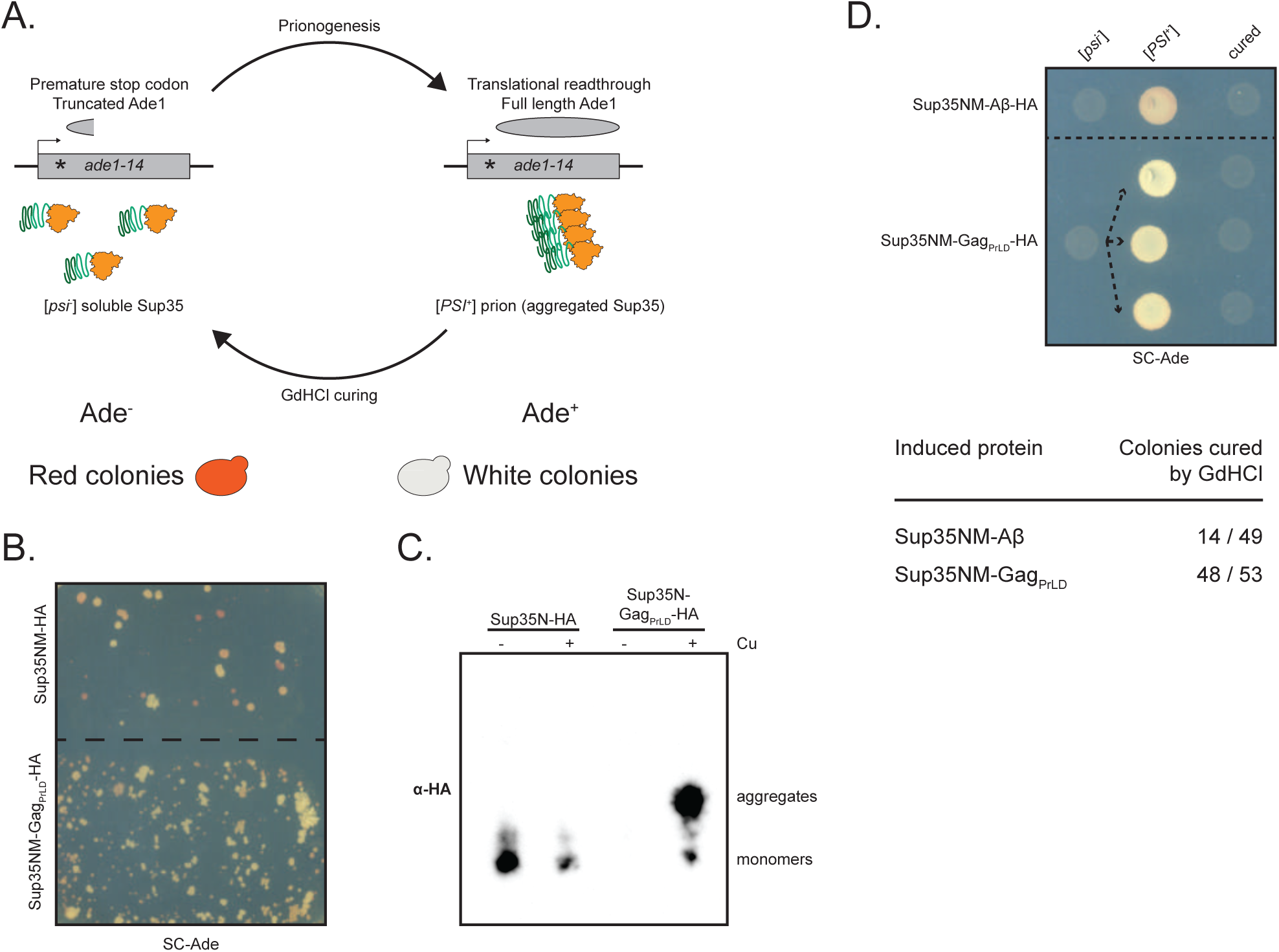
Gag_PrLD_ nucleates a Sup35-based prion reporter. (*A*) Schematic of the prionogenesis assay using the *ade1-14* allele containing a premature stop codon. Soluble Sup35 terminates translation at the premature stop codon, yielding a non-functional, truncated Ade1 (N-succinyl-5-aminoimidazole-4-carboxamide ribotide synthetase); yeast cannot grow on media lacking adenine (SC-Ade) and a red pigment develops. Sup35 aggregated into the prion state allows for translational readthrough and production of functional Ade1; yeast grow on SC-Ade and appear white. (*B*) Qualitative prionogenesis of Sup35NM fusions; growth on SC-Ade indicates either a suppressor mutation or [*PSI^+^*] prionogenesis. Expression of Sup35 fusions were induced with 150 μM CuSO_4_. A representative image of at least 3 experiments is shown. (*C*) SDD-AGE analysis of Sup35N-HA with and without Gag_PrLD_ fusion. Expression of Sup35 fusions were induced with 100 μM CuSO_4_. Monomers and high-molecular weight aggregates of chimeric proteins were detected with anti-HA antibody. A representative image of at least 3 experiments is shown. (*D*) Curing of Ade^+^ colonies by guanidine hydrochloride (GdHCl) of Sup35NM-HA chimeras. One [*psi^-^*] Sup35NM-Aβ fusion control strain is shown induced to [*PSI^+^*] and cured. Three independent inductions of a [*psi^-^*] Sup35NM-Gag_PrLD_ fusion are shown induced to [*PSI^+^*] and cured. [*PSI^+^*] yeast grow on SC-Ade while [*psi^-^*] and cured yeast do not. The table below shows the guanidine curability of Ade^+^ colonies induced by chimeric constructs.

### The Gag_PrLD_ is required for Ty1 transposition

Given that the Gag_PrLD_ promotes prionogenesis of a Sup35-based reporter, we investigated its functional role in Ty1 transposition. In a *Saccharomyces paradoxus* strain lacking genomic Ty1 elements (57, 58), we first deleted the PrLD from Gag in a Ty1 element provided on a plasmid and tagged with the robust and sensitive *his3-AI* retrotranscript indicator gene (59) (Fig. 3*A*). This marker contains a mutant *his3* gene split by an antisense artificial intron (AI) that is inserted at the 3’ untranslated region of Ty1 in the opposite transcriptional orientation. The AI is in the correct orientation to be spliced only in Ty1*his3-AI* RNA; cDNA reverse transcribed from this product results in a functional *HIS3* allele. Insertion into the genome, either by integration or recombination, allows cells to grow on media lacking histidine. The frequency of His^+^ prototrophy is a direct measure of Ty1*his3-AI* retrotransposition or cDNA recombination, collectively known as retromobility. Deletion of the Gag_PrLD_ in a complete Ty1*his3-AI* element overexpressed under the *GAL1* promoter completely abolished retromobility (Fig. S4*A*), despite retaining normal Gag protein levels (Fig. S4*B*). However, the PrLD region of Gag contains *cis*-acting RNA signals required for efficient reverse transcription (60, 61). To distinguish between a functional role in retrotransposition of the PrLD in the Gag protein versus the role of the RNA sequences that encode for the PrLD, we used a two-plasmid system to separate Ty1 RNA and protein functions (Fig. 3*B*). A helper-Ty1 encodes a functional mRNA, providing protein products, but lacks a 3’ LTR thus disrupting *cis*-acting signals required for reverse transcription. Mini-Ty1*his3-AI* lacks complete open-reading frames (ORFs) but contains *cis*-acting signals for dimerization, packaging, and reverse transcription of mini-Ty1*his3-AI* RNA (61, 62). Retromobility is monitored through the *his3-AI* reporter. In the two-plasmid assay, deletion of the Gag_PrLD_ also inhibits retromobility (Fig. 3*C-D*), despite producing normal levels of Gag protein (Fig. 3*E*), confirming a critical contribution from the PrLD in the Gag protein to retromobility.

**Fig 3.**
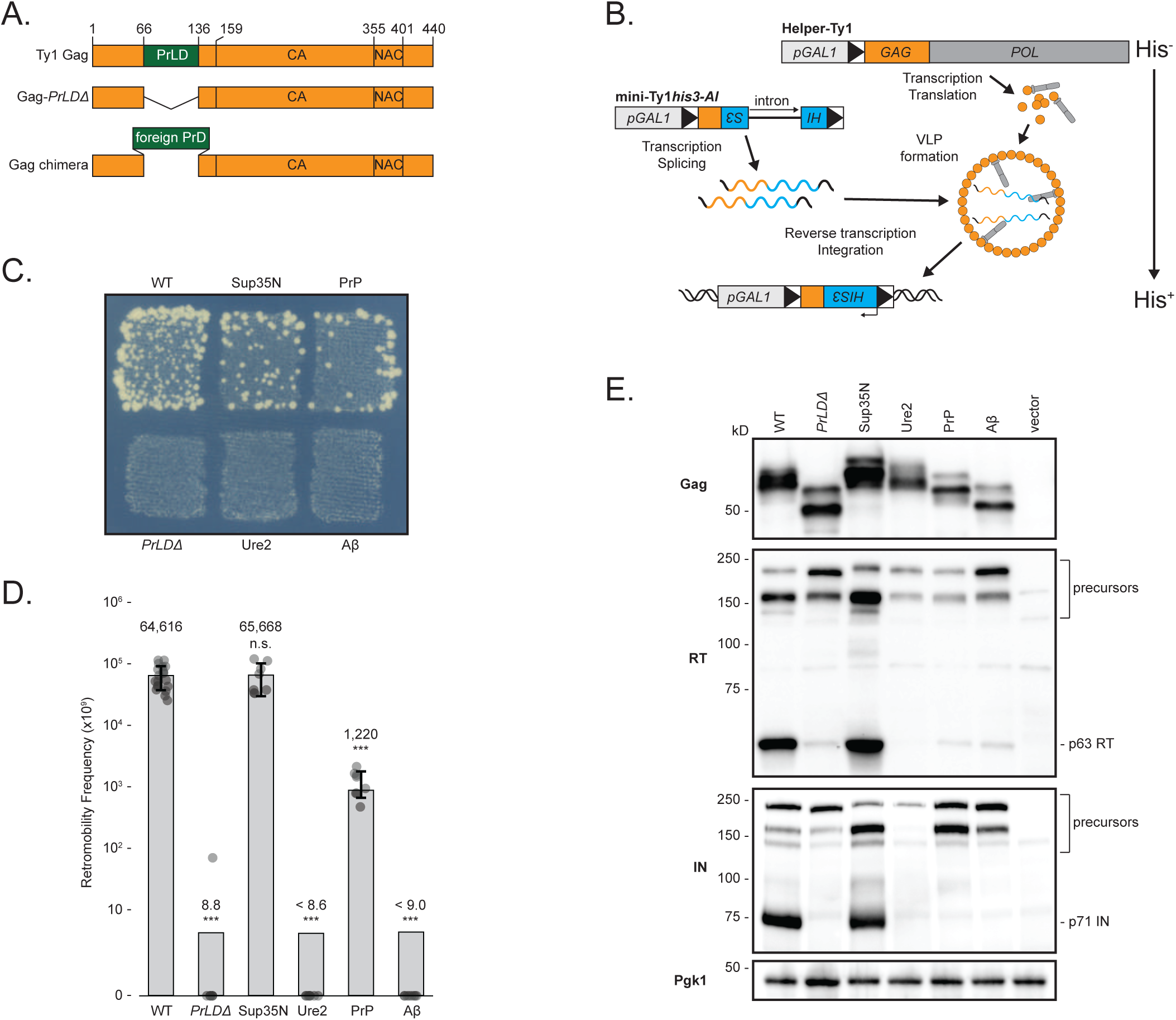
Ty1 Gag chimeras containing known PrDs produce stable Gag but have a range of transposition and proteolytic maturation phenotypes. (*A*) Schematic of Ty1 Gag constructs. The Ty1 Gag PrLD is intact in wildtype (WT), deleted in *PrLDΔ*, and replaced with known PrDs in the chimeras. (*B*) Schematic illustrating the two-plasmid system separating Ty1 RNA and protein functions. Helper-Ty1 encodes a functional mRNA, providing protein products, but lacks a 3’ LTR thus disrupting *cis*-acting signals required for reverse transcription. Mini-Ty1*his3-AI* lacks complete ORFs but contains *cis*-acting signals for dimerization, packaging, and reverse transcription of mini-Ty1*his3-AI* RNA. The *his3-AI* indicator gene detects retromobility of mini-Ty1*HIS3* cDNA. (*C*) Qualitative retromobility of chimeric Gag constructs in the two-plasmid system. Colony growth on a medium lacking histidine indicates a retromobility event. A representative image of at least 3 replicates is shown. (*D*) Quantitative mobility assay of galactose-induced cells. Each bar represents the mean of at least eight independent measurements, displayed as points, and the error bar ± the standard deviation. Error bars are omitted for *PrLDΔ*, Ure2, and Aβ chimeras that did not transpose; one retromobility event was observed in one replicate of *PrLDΔ*. Adjusted retromobility frequency is indicated above the bars. For Ure2 and Aβ, frequencies are indicated as less than the calculated frequency if one retromobility event had been observed. Significance is calculated from a two-sided Student’s *t*-test compared with WT (n.s. not significant, \*\*\**p* < 0.001. Exact *p*-values are provided in Supplementary Table 1). (*E*) Protein extracts prepared from galactose-induced cells expressing the indicated Gag constructs in the two-plasmid system were immunoblotted for the protein indicated on left. Polypeptide precursors are bracketed and mature RT and IN sizes are noted on right. Pgk1 serves as a loading control. Migration of molecular weight standards is shown alongside the immunoblots. A representative image of at least 3 replicates is shown.

### Ty1 mobility of Gag chimeras containing foreign PrLDs

To better understand the nature of the PrLD’s contribution to retromobility, we asked whether the Gag_PrLD_ sequence is uniquely capable of facilitating retromobility. Since the Gag_PrLD_ has prionogenic properties and sequence similarity to prions, we created chimeric Ty1 Gags in which the PrLD is replaced with prion domains from well-studied prions and aggregating proteins (Fig. 3*A*). We chose the yeast prions Sup35 and Ure2, the mouse prion protein PrP, and the Alzheimer’s disease-associated human Aβ_1-42_ using domains predicted computationally (Fig. 1) (52, 53, 63). Chimeric Ty1 elements on the helper-Ty1 plasmid were co-expressed with mini-Ty1*his3-AI*, and the level of Ty1 mobility was determined. Remarkably, substitution of the Gag_PrLD_ with the prion domain from yeast Sup35 or mouse PrP supported Ty1 retromobility in qualitative (Fig. 3*C*) and quantitative retromobility assays (Fig. 3*D*). Gag_Sup35N_ retromobility is not significantly different from wildtype, whereas Gag_PrP_ is an order of magnitude lower, although still readily detectable on a qualitative plate assay. Replacing the PrLD sequence disrupts RNA signals which is reflected in the single plasmid assay, in which Gag_Sup35N_ and Gag_PrP_ chimeras have dramatically reduced retromobility (Fig. S4*A*), despite producing similar Gag protein levels (Fig. S4*B*), highlighting the importance of separating protein and RNA function with the two-plasmid assay.

Retromobility measured as the frequency of His^+^ prototrophs formed from *his3-AI* tagged elements includes both new chromosomal integrations likely created via retrotransposition, and recombination of the spliced cDNA with homologous sequences present on the mini-Ty1*his3-AI* plasmid. To assess whether the chimeras support retrotransposition or merely recombination, we distinguished the two by monitoring histidine prototrophy after segregating the helper and mini-Ty1*his3-AI* plasmids (Fig. S4*C*). In our strain background with the wildtype two-plasmid system, 4% of retromobility events were due to recombination with either of the plasmids. The Gag_Sup35N_ and Gag_PrP_ chimeras had modestly increased recombination events, although only Gag_Sup35N_ reached statistical significance (*p*=0.024) (Fig. S4*D*). We conclude that the Gag chimeras support *de novo* retrotransposition and cDNA recombination remains a minor pathway (64, 65).

### Effect of Gag_PrLD_ chimeras on Ty1 protein level and maturation

The result that the Gag_PrLD_ can be replaced by foreign prion sequences indicates its function is not unique to the PrLD sequence and may be the same as provided in aggregation-prone proteins. However, not all the disordered domains tested in Gag chimeras supported transposition. Ty1 chimeras containing the domains from yeast Ure2 or human Aβ did not transpose (Fig. 3*C-D*). All the chimeric Gags were expressed at similar levels (Fig. 3*E*), arguing against different transposition phenotypes due to effects on protein stability from the foreign prion domains. The substituted prion domains are of various sizes, and Gag chimeras had predicted electrophoretic mobilities. Gag proteolytically matures from p49 to p45 and is subject to post-translational modifications, often resulting in multiple bands observed by western blot (3). To determine whether the Gag chimeras affected protein maturation, we assessed the relative levels of mature RT and IN by western blotting with antibodies specific to each protein. Deletion of the PrLD results in dramatically reduced mature RT and IN levels (Fig. 3*E*). The Gag_Sup35N_ chimera transposed as well as wildtype and produced equivalent levels of mature RT and IN. The transposition-deficient chimeras, Gag_Ure2_ and Gag_Aβ_, have very reduced levels, comparable to Gag*_PrLDΔ_*. Interestingly, Gag_PrP_ supports transposition, although reduced from wildtype, and has low levels of mature RT and IN. These results raise the possibility that Gag chimeras can block PR function and production of mature RT and IN that are essential for Ty1 mobility.

### Ty1 Gag*_PrLDΔ_* and Gag chimeras fused to GFP affect aggregation and localization

Proteolytic maturation of RT and IN via PR occurs within VLPs (66), which are believed to be assembled in retrosomes (20–22). Gag fused to green fluorescent protein (GFP) has been used as a reporter for retrosome assembly and location (67), therefore we examined formation of cytoplasmic foci of wildtype, mutant, and chimeric Gag-GFP in the Ty-less background. Wildtype Gag-GFP fusions formed discrete cytoplasmic foci, as previously reported using this construct, but deleting the PrLD resulted in diffuse localization throughout the cytoplasm (Fig. 4). We found that a 24 hr galactose induction, shorter than 48 hr-induction used above, was ideal for live-cell microscopy and GFP-detection as yeast cultures are in log-phase growth (Fig. S5). 24 hr induced Gag_Sup35N_ formed similarly discrete foci patterns as wildtype Gag, whereas Gag_Ure2_ had diffuse localization similar to Gag*_PrLDΔ_*. Gag_PrP_ supports transposition and predominately formed foci similar to wildtype, but also had a modest fraction of cells containing a visually distinct fluorescent morphology that appears as a single, large, very bright focus. Ty1 Gag_Aβ_ does not transpose, yet formed foci and an even larger fraction of cells contained these single, large foci. Forming Gag-GFP foci correlates with a requirement for transposition but, as Gag_Aβ_ shows, is not sufficient.

**Fig 4.**
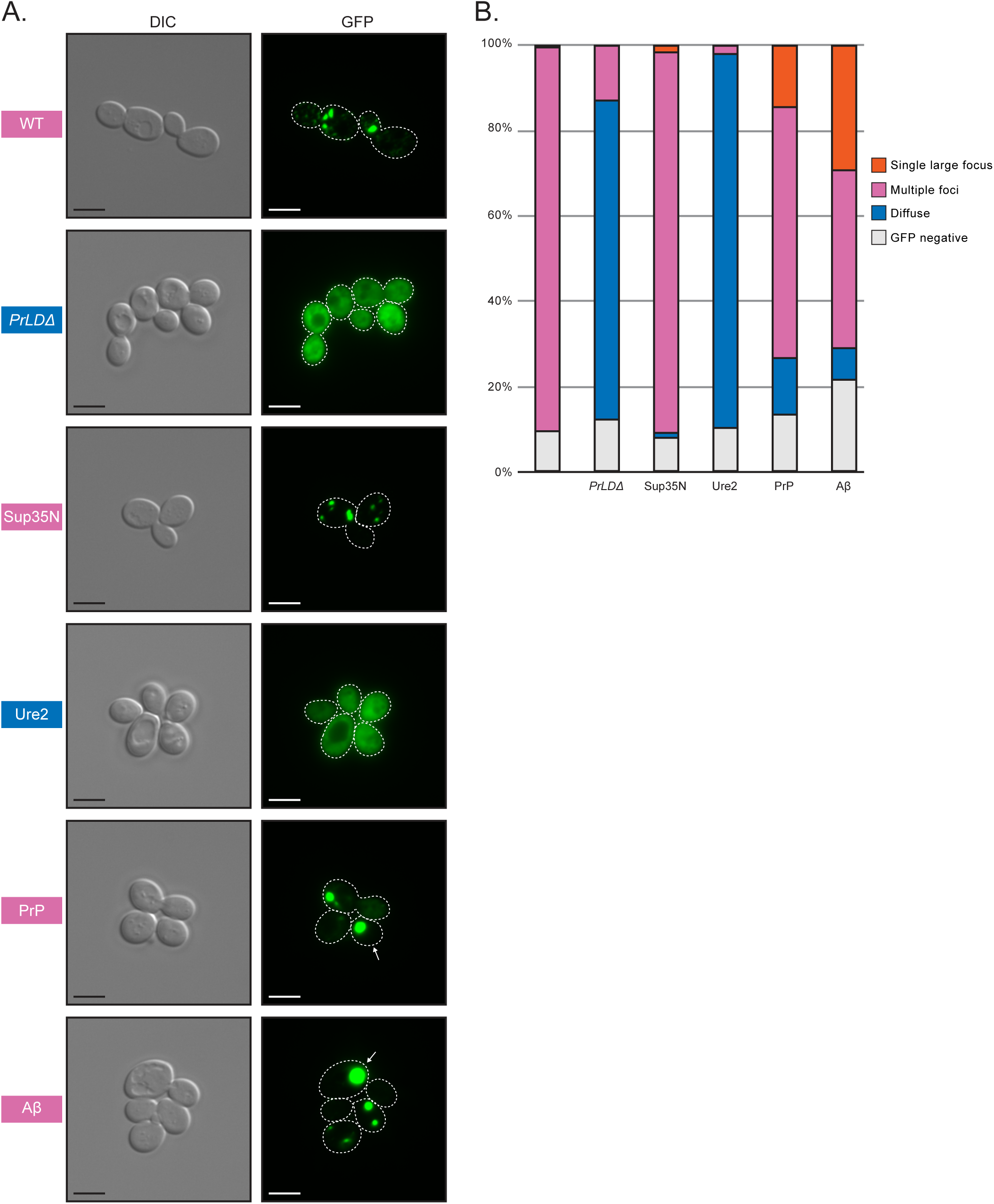
Foci detected in cells expressing wildtype Gag, the Gag*PrLDΔ* mutant, and Gag-PrLD chimeras fused to GFP. (*A*) Live-cell yeast fluorescence microscopy of strains expressing chimeric Gag-GFP after 24 hr galactose induction. Normaski (DIC) and GFP channels are shown with cell outlines added to GFP channels based on DIC images. The strain labels are colored to match the most common foci observed. White arrows indicate cells with a single large focus. Scale bars represent 5 μm. (*B*) Quantitation of categories of foci observed as a percentage in at least 300 cells. The multiple foci category includes cells with multiple large foci, one or more small foci, or a combination of both sizes. Cell counts are provided in Supplementary Table 2.

In addition, we investigated the structures formed by Gag-GFP chimeras in fixed yeast cells by thin section transmission electron microscopy (TEM) (Fig. S6) using methods similar to those used for detecting Ty1 VLPs (22). Wildtype Gag-GFP produced electron-dense structures that appear similar to VLPs but look incomplete or incorrectly assembled, lacking a circular shell with a hollow interior. Gag*_PrLDΔ_* did not form any VLP-like structures detectable in micrographs. The Gag_Sup35N_ strain produced tubular or filamentous structures, also not resembling proper VLPs. And strikingly, the Gag_Aβ_ strain formed large densities in defined regions of the cell, instead of clusters of particles or filaments across the cytoplasm, perhaps corresponding to the single large foci seen by fluorescent microscopy. These results suggest that Gag-GFP can reveal severe assembly defects as evidenced by Gag*_PrLDΔ_* but GFP may confer aberrant VLP assembly properties when wildtype or chimeric Gag-GFP fusions are produced in cells.

### The Ty1 Gag*_PrLDΔ_* and Gag chimeras affect VLP assembly

To evaluate VLP assembly in the chimeras using the two-plasmid system, we examined Gag sedimentation profiles of yeast lysate run through a 7-47% continuous sucrose gradient, as previously reported (15, 58, 68). Wildtype VLPs accumulated in more dense sucrose fractions near the bottom half of the gradient, with peak fractions indicated by a bar (Fig. 5). Gag*_PrLDΔ_* appears unable to assemble complete VLPs, as Gag in these mutants accumulated in less dense sucrose fractions near the top of the gradient. The transposition-competent chimeras Gag_Sup35N_ and Gag_PrP_ had similar sedimentation profiles as wildtype, whereas transposition-deficient Gag_Ure2_ accumulated near the top of the gradient like Gag*_PrLDΔ_*. Gag_Aβ_ does not support retrotransposition, but peaked in similar fractions as wildtype, although somewhat more broadly distributed across the gradient.

**Fig 5.**
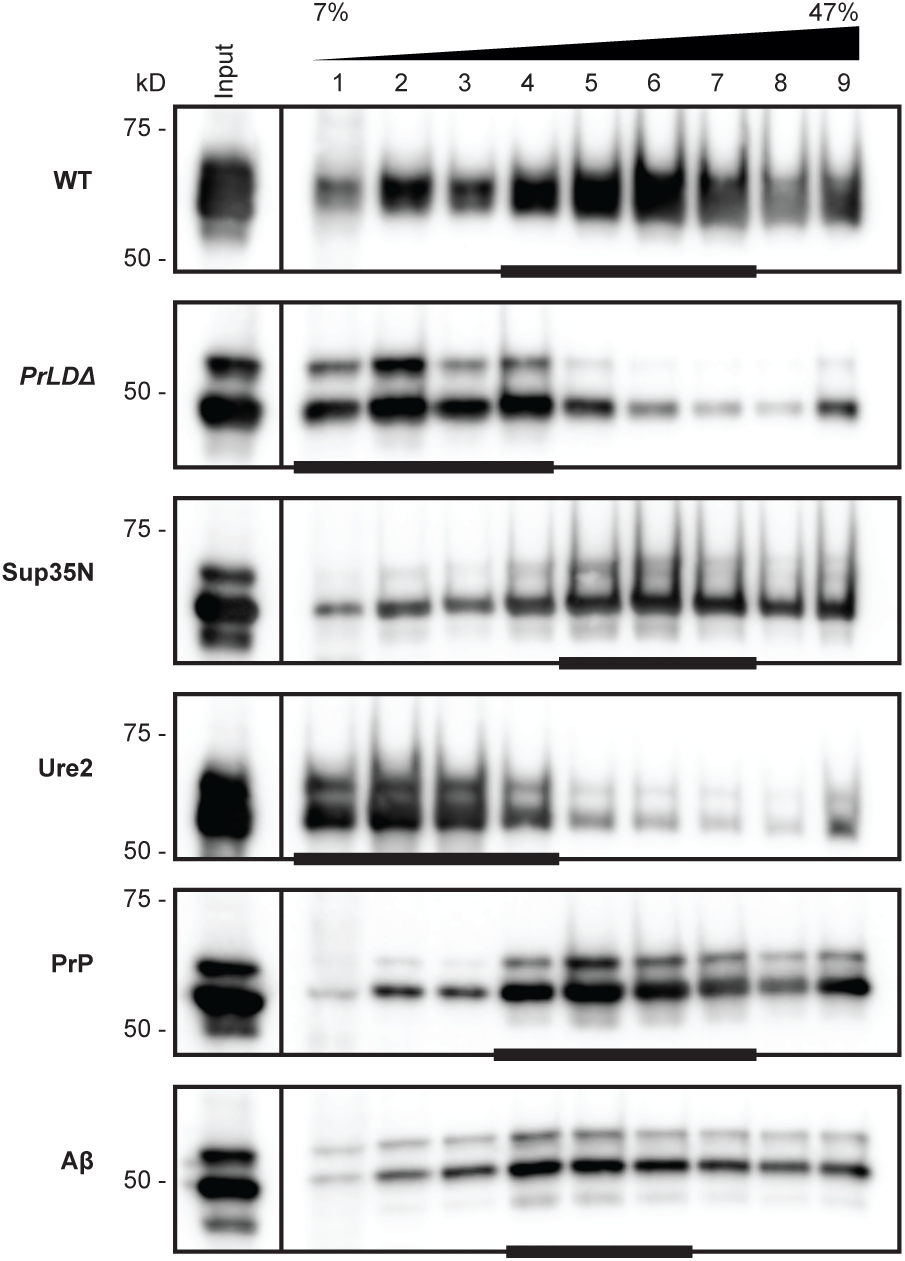
Transposition-incompetent Gag chimeras disrupt VLP assembly. Protein extracts from galactose-induced yeast cells (Input) were fractionated over a 7–47% continuous sucrose gradient and immunoblotted for Gag. Expression plasmids and molecular weight standards are noted alongside the blots. The bars at the bottom of blots denote peak Gag fractions containing more than 1/9 of the Gag signal across the gradient, as determined by densitometric analysis. A representative image of at least 3 replicates is shown.

To further examine the VLPs assembled by each chimera, we visualized thin sections of fixed yeast cells by TEM. Cells overexpressing the wildtype two-plasmid Ty1 system produced large clusters of VLPs (Fig. 6). VLPs are characteristically round with an electron dense shell and their interior appears hollow in micrographs. Importantly, these particles were not observed in the parental yeast strain expressing empty vectors. Ty1 VLPs are heterogeneously sized and are approximately 30-80 nm in diameter, based on previous measurements of purified particles (69, 70). In thin section TEM, particles may be in different Z-planes when sectioned, therefore masking the diameter of a roughly spherical particle, and preventing quantitative particle size data collection from thin section TEM. With this limitation in mind, we measured particle diameters from several cells in multiple micrographs to estimate an approximate size, and found wildtype particles ranging from 40-80 nm, with a median diameter of 59 nm (Fig. S7), largely agreeing with previous reports of purified particles.

**Fig 6.**
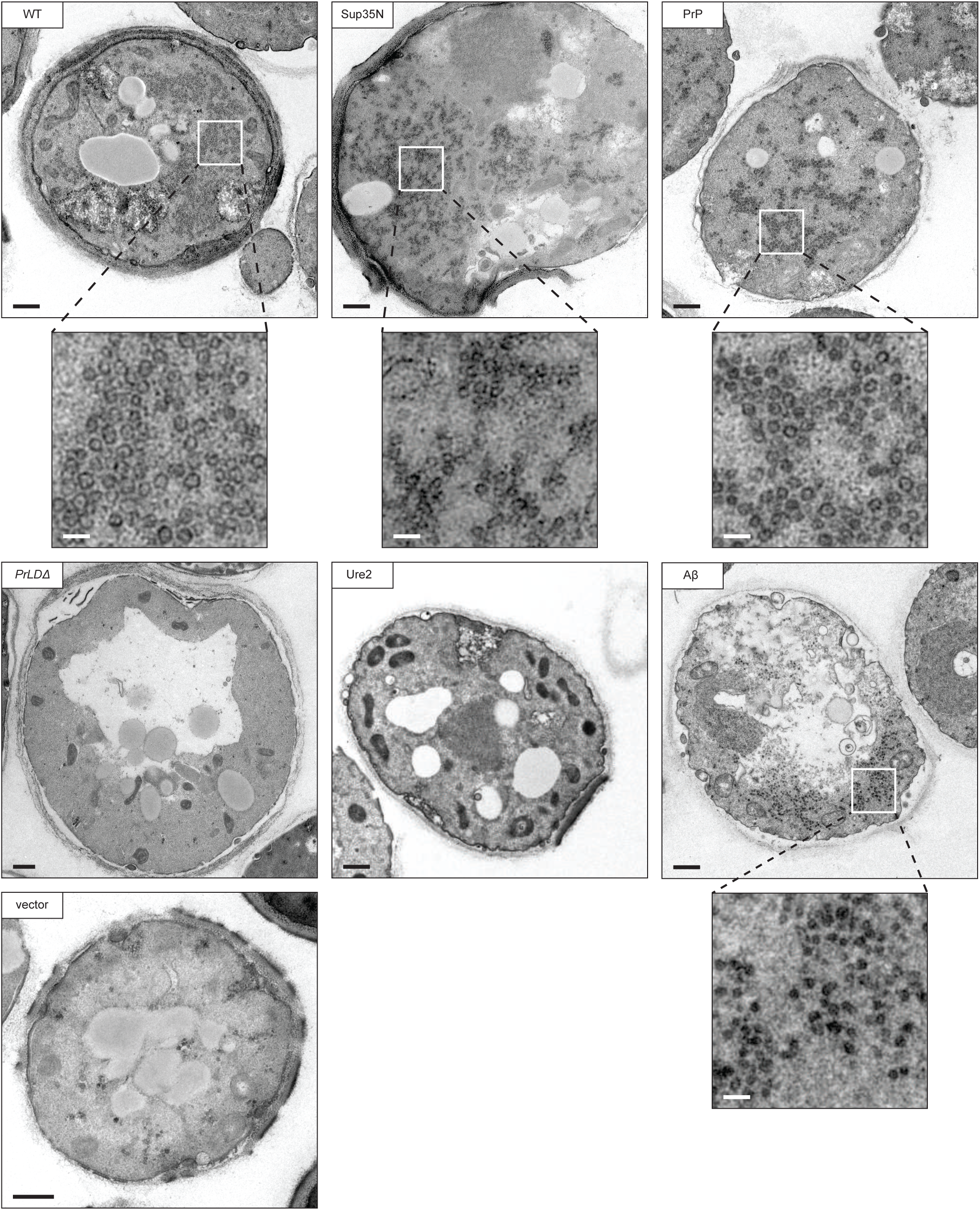
Transposition competent Gag chimeras support VLP production. Thin-section TEM of galactose-induced cells expressing Gag chimeras. Representative cells are shown, those containing VLP clusters include zoomed in cutouts to highlight VLPs. The black bars represent 500 nm, the white bars represent 100 nm.

We did not observe any cells producing VLPs in the Gag*_PrLDΔ_* mutant, in agreement with Gag*_PrLDΔ_*-GFP imaging and sucrose sedimentation profiles. Taken together, these data lead us to conclude that the PrLD is required for Ty1 VLP assembly. The transposition-deficient Gag_Ure2_ chimera also did not assemble VLPs as monitored by thin section TEM, again agreeing with sucrose sedimentation results. The two transposition-competent chimeras, Gag_Sup35N_ and Gag_PrP_, assembled VLPs similar in size and appearance to wildtype. These chimeras also produced large numbers of particles in each cell, although consistently appearing somewhat more dispersed throughout the cell than wildtype particle clusters. Interestingly, Gag_Aβ_ does not support retrotransposition, but has a similar sucrose sedimentation profile as wildtype, suggesting it may assemble particles that are defective for transposition. In thin section TEM, we observed particles in cells expressing Gag_Aβ_ that are visually distinct from wildtype. The most striking difference is that these particles do not have the characteristic hollow center and instead appear electron-dense throughout. They are smaller than wildtype with a median diameter of 42 nm (Fig. S7), and, like the Gag_Sup35N_ and Gag_PrP_ chimeras, are produced in large numbers of particles but are dispersed throughout the cell. Together, these results illustrate the robust and flexible nature of VLP assembly. However, our data also underscore the requirement for PrLD functionality as yeast and mammalian Gag-prionogenic chimeras form VLPs *in vivo* whereas the Gag*_PrLDΔ_* mutant does not.

## Discussion

The data presented here permit several conclusions about prionogenic domains, the functional organization of Ty1 Gag, and VLP assembly. Our results demonstrate that the Ty1 Gag protein contains a novel prion-like domain that is required for VLP assembly and retrotransposition. The Gag_PrLD_ has intrinsic prionogenic properties demonstrated by a cell-based Sup35 reporter assay, and its function in Ty1 transposition can be replaced by certain yeast and mammalian bona fide prion domains. Our findings also raise interesting unresolved questions about sequence constraints of PrLDs and how widespread PrLD functions are across retroelements. Finally, this work suggests using Ty1 as an *in vivo* screening platform to study intrinsically disordered domains.

### Prion properties of the Ty1 Gag_PrLD_

We have examined prionogenic properties of the Ty1 Gag_PrLD_ using an established cell-based assay in which the newly discovered Gag_PrLD_ is fused to the N- and NM-domains of Sup35. Nonsense readthrough is measured by auxotrophic growth and colony color, aggregate formation is monitored biochemically with SDD-AGE, and curability is assessed after GdHCl treatment. It will be informative to further characterize prionogenic properties of the Ty1 Gag_PrLD_ using additional assays on various Gag_PrLD_ fusion constructs, including fused to Sup35C, and measuring binding of the amyloid-sensitive dye thioflavin-T, non-Mendelian inheritance, aggregation in SDD-AGE, and GFP localization patterns (71, 72).

### RNA-contributions to Gag_PrLD_ function

To disentangle protein-level effects of mutations in the Ty1 Gag PrLD from mutations of *cis*-acting RNA sequences, we separated RNA and protein function in a two-plasmid system. Retromobility is considerably lower in the two-plasmid system (Fig. 3C) than a single-plasmid expressing the intact transposon (Fig. S4A). Nonetheless, the two-plasmid system provides a wide dynamic range allowing for sensitive measurement of the impact of Gag_PrLD_ chimeras on retromobility. The Gag_Sup35N_ chimera restored retromobility in the two-plasmid system but had a severe retromobility defect in the single-plasmid assay. Based on our current knowledge of functional contributions from Ty1 *cis*-acting RNA sequences to retrotransposition, this is likely due to disruption of the pseudoknot sequences in the RNA region that encodes for the PrLD (60, 61). It may be possible to engineer an equivalent pseudoknot sequence in the Sup35N-encoding RNA and restore retromobility in a single-plasmid Gag_Sup35N_ chimera. Our chimeric proteins may provide a useful platform to interrogate RNA requirements.

### Sequence requirements of the Ty1 Gag_PrLD_

We replaced the Gag_PrLD_ with exogenous prion domains selected based on computational predictions and the literature. We chose the entirety of Aβ_1-42_ and the complete N-terminal domain of Sup35_2-123_. We introduced the highest scoring 60 amino acid stretch predicted by PLAAC, Ure2_17-76_, which is within the established prion domain reported as the first 89 amino acids (52, 63). The infectious PrP 27-30 isoform initially isolated is roughly 142 amino acids long and spans from approximately residues 90 to 230 (73), but shorter truncations still display prion phenotypes (74–76) and PrP_90-159_ is sufficient to induce prionogenesis in a yeast-based assay (53). The PrP_121-231_ fragment is soluble, and its structure has been determined using solution NMR (49, 77). We introduced PrP_90-159_ as a Ty1 Gag chimera based on prior success in yeast. It will be interesting to examine other regions of PrP for function when present in Gag.

The sequence features constraining Ty1 PrLD function are not yet well-defined. Intriguingly, both the transposition-competent Gag chimeras (Sup35 and PrP) are from proteins with oligopeptide repeats associated with prionogenesis (78, 79). However, the PrP sequence introduced as a Ty1 Gag chimera in this study does not contain these repeats. Moreover, the Ty1 Gag_PrLD_ does not have equivalent repeats of 8-10 amino acids. Instead, like other reported prion domains, the Gag_PrLD_ is Q/N-rich and is depleted of charged residues. Additionally, a large number of prolines in the Gag_PrLD_ likely prevents secondary structure formation and starkly contrasts with the highly alpha-helical folding of the Gag capsid domain (15, 68). Further investigation will be required to understand the sequence parameters, such as length, amino acid composition, charge, or oligorepeats, that govern function of the Gag_PrLD_. Transposition-deficient Gag_PrLD_ chimeras may be analyzed by reversion analysis to select mutations that restore transposition. Characterizing the revertants could reveal incompatibility with PR or other Pol proteins, rather than early VLP assembly steps.

### Interaction of the Gag_PrLD_ with the host and environment

Many host genes have been identified that activate or restrict Ty1 transposition (80–83), and examining genetic interactions with the Gag_PrLD_ may provide regulatory insights. Prion domains have been proposed to be protein-specific stress sensors that allow cells to respond to environmental conditions (84–86). Substituting the Gag PrLD may therefore change Ty1 regulation and create genetic interaction partners selective for specific Gag chimeras. For example, modulators of Sup35 prionogenesis could specifically modulate Gag_Sup35N_ but not wildtype Ty1. Conversely, Gag chimeras may no longer be subject to regulation by Ty1 modulators.

### Gag chimeras reveal varied deficiencies across the Ty1 life cycle

Different Gag_PrLD_ chimeras had different phenotypes across the Ty1 life cycle. Gag_PrP_ supported retromobility, although less well than wildtype or Gag_Sup35N_. Whereas Gag_PrP_ produces VLPs that appear to have wildtype morphology by TEM (Fig. 6) and have similar sedimentation profiles to wildtype (Fig. 5), Gag_PrP_ accumulates low levels of mature RT and IN (Fig. 3E). This could indicate an incompatibility of Gag_PrP_ as a substrate for PR. Another possibility is that Gag_PrP_ VLPs inefficiently incorporate Gag-Pol or are partially defective in ways not detectable by TEM or sedimentation. The reduced retromobility of Gag_PrP_ may be explained by impaired RT and IN protein maturation. Meanwhile, Gag_Aβ_ produces particles that do not support retrotransposition or Pol maturation. These particles lack the characteristic hollow center observed in TEM of wildtype VLPs (Fig. 6) and are noticeably smaller in diameter (Fig. S7). These observations highlight that VLP assembly is robust but underscores the point that simply assembling particles is not sufficient for transposition and that assembling correct VLPs is required for proteolytic maturation. It will be informative to measure packaging of the mini-Ty1 RNA into chimeric VLPs. Ultimately, cDNA synthesis requires both the mature enzymes and the RNA substrate to be present in VLPs. Our sedimentation and TEM results presented here build upon previously published sedimentation experiments (15, 58, 68), and strengthen the value of sedimentation as a proxy for VLP assembly. Nonetheless, the value of TEM is exemplified by the Gag_Aβ_ chimera, which sediments similarly to wildtype but TEM reveals aberrant particle morphology.

Gag-GFP fusions have been used as a proxy for Ty1 retrosomes (67), although we have not formally tested for Ty1 RNA co-localization in our specific system. We used a previously published wildtype GFP-fusion construct that contains the mature Gag (p45) and not a full-length element. The utility of Gag-GFP is shown by the cellular mislocalization observed in Gag chimeras. However, GFP is a 26 kD protein and fusion impaired proper VLP formation (Fig. S6), perhaps interfering with Gag-Gag contacts that must be made to assemble the complete particle structure. Examining the PrLD fused to GFP alone, without the full Gag protein, or testing a Gag truncation that lacks the NAC domain, will indicate the minimal region that promotes foci formation and if RNA recruitment is required. Ty1 Gag contains a NAC and binds Ty1 RNA, but also binds diverse RNAs *in vitro* and cellular mRNAs associate with Ty1 VLPs (16, 17, 62, 87– 89). Whether Ty1 RNA, specifically, is required to form foci or to nucleate VLP assembly, or if there is an RNA requirement at all, will require further study. It remains to be determined whether the GFP foci and VLP nucleation site is associated with any subcellular locales, as has been previously proposed at the endoplasmic reticulum (67).

The Gag-GFP foci may mature over time, as an increased percentage of Gag*_PrLDΔ_* cells had foci after 48 hr of induction compared to 24 hr. These foci are proxies for retrosomes and therefore represent an early step of the Ty1 life cycle preceding VLP assembly, protein maturation, and transposition. After 48 hr of galactose induction, cultures enter stationary-phase growth and have higher levels of GFP-negative cells which often appear to have a wrinkled, potentially senescent morphology (Fig. S5). We, therefore, chose to examine Gag chimeras by fluorescent microscopy after 24 hr, but our pilot experiments at 48 hr anecdotally suggested more bright, single large foci. This observation would be consistent with a kinetic component to retrosome formation and may represent a progression from LLPS towards hydrogel formation. Whether such a gel would be an irreversible phase that is unable to dissolve and proceed with VLP formation remains to be determined.

### Does the Ty1 retrosome constitute a phase-separated compartment?

Wildtype cells assemble discrete VLPs that can be found throughout the cell but are often observed in a particular region of the cytoplasm, and even the wildtype Gag-GFP assembled discrete structures, observed by TEM. However, the Gag_Aβ_-GFP strain produced large densities that may correspond to large foci observed by fluorescence microscopy. These assemblies would be consistent with LLPS compartments containing high concentrations of Gag-GFP that stall and cannot complete VLP assembly; however, we have not examined LLPS properties such as concentration-dependence, droplet merging, or internal mixing (23). Prion-like domains can drive formation of a gradient of assemblies, from LLPS to hydrogels and amyloid-like fibers. The Ty1 Gag chimeras may exhibit a spectrum of these morphologies. The filamentous assemblies formed by Gag_Sup35N_-GFP are potentially similar to Sup35 amyloid fibers observed *in vitro*, and Gag_Aβ_-GFP may form liquid droplets. Sup35, while canonically known for its ability to form amyloid fibers as a prion, has more recently been appreciated to undergo LLPS upon a decrease in cytosolic pH and can mature over time into a gel-like condensate (85, 90). Whereas wildtype Gag allows for VLP assembly to proceed and supports transposition, perhaps transiently existing in an LLPS state, chimeras may become blocked along the retrosome and VLP assembly pathway, resulting in the striking structures observed by fluorescence microscopy and TEM. Further work will be required for the rigorous characterization necessary to declare the Ty1 retrosome or other assemblies formed by Gag chimeras an LLPS compartment. Ty1 provides a promising system to unite studies of prion and LLPS pathways.

### An interchangeable platform to study PrLD and LLPS domains in living cells

The condensate-forming property, but not the prion-forming property, of Sup35 is conserved across 400 million years from *S. cerevisiae* to *Schizosaccharomyces pombe*, emphasizing the evolutionary importance of this ancient phenotype (85). Our discovery of the Ty1 PrLD raises the possibility that LLPS may, too, be widespread among retroelements. Our preliminary computational analyses of Pseudoviridae (Ty1/copia) retroelement family members reveal predicted PrLDs in not only the closely-related yeast Ty2, but also in distantly-related plants in the *Oryza* element *Retrofit* and the *Arabidopsis* elements *Evelknievel* and AtRE1. The human retrotransposon LINE-1 phase separates and retrotransposition is associated with cancer (91) and age-associated inflammation (92, 93). A condensate-hardening drug was found to block human respiratory syncytial virus replication which occurs in virus-induced inclusion bodies (94), highlighting the potential of the Ty1 platform to contribute to new anti-viral and other human-health therapeutics. The Ty1 Gag chimera strategy developed here may prove to be a useful platform to study prion-like and LLPS-forming domains due to the genetic tractability of yeast and the suite of robust and sensitive *in vivo* assays developed for Ty1.

## Materials and Methods

### Bioinformatic analyses

PLAAC (http://plaac.wi.mit.edu/) (42) was used with default settings: core length of 60 and 100% *S. cerevisiae* background probabilities. ArchCandy (https://bioinfo.crbm.cnrs.fr/index.php?route=tools&tool=7) (43) was used with a score threshold of 0.500 and the transmembrane regions filter off; the sum of scores data is presented. PrDOS (https://prdos.hgc.jp/) (44) was used with the default 5% FDR and the disordered probability threshold set to 0.5. GlobPlot2.3 (http://globplot.embl.de/) (45) was used with default settings with Russell/Linding propensities. Bioinformatic outputs were uniformly plotted using a custom script using the base plot() and rect() functions in R version 3.5.2. Structure analysis was performed using PyMOL v1.5.0.5 with the “align” command.

### Yeast strains and media

Yeast strains with full genotypes are listed in Supplementary Table 3. Standard yeast genetic and microbiological techniques were used in this work (95). Prion nucleation experiments were performed in GT409, an *S. cerevisiae* strain that is [*psi^-^ pin^-^*] and harbors the *ade1-14* allele which contains a premature stop codon (kindly provided by Y. Chernoff) (53). Ty1 assays were performed in the DG3582 background, a Ty-less *S. paradoxus* derivative of DG1768 (57, 58). For galactose induction in liquid media, starter cultures were grown overnight at 30 °C in synthetic complete (SC) dropout media containing 2% raffinose, diluted 1:20 into media containing 2% galactose, and grown at 22 °C for 48 hours.

### Plasmids and cloning

Plasmids, primers, and gene fragments are listed in Supplementary Tables 4-6. Detailed descriptions of plasmids and cloning are provided in SI Materials and Methods.

### Prion nucleation and curing

[*PSI+*] induction was assayed in a [*psi-*] strain for chimeric plasmids under a P*_CUP1_* promoter; yeast cells were grown at 30 °C. Yeast were grown on SC-Ura for 2 days, replica plated to SC-Ura ± 150 μM CuSO_4_ and grown for 2 days, then replica plated to SC-Ade and grown for approximately 10 days until imaged. Following prion nucleation, Ade^+^ colonies were cured of [*PSI^+^*] by guanidine hydrochloride (GdHCl). First, the induction plasmid was counter selected on FOA and single colonies were isolated. Then, Ade^+^/Ura^-^ colonies were passaged as single colonies on YPD spotted with 10 or 25 μL of 5 M GdHCl until red-pigmented colonies developed.

### SDD-AGE

Semi-denaturing detergent-agarose gel electrophoresis (SDD-AGE) was adapted from published methods (53, 54). Detailed protocols are described in SI Materials and Methods.

### Ty1*his3-AI* mobility assays

Ty1 retromobility events were detected using the *his3-AI* retromobility indicator gene (59) by qualitative and quantitative assays as previously described (15, 58). Detailed protocols are described in SI Materials and Methods.

### Immunoblotting

Total yeast protein was prepared by trichloroacetic acid precipitation and immunoblotted using standard techniques (58, 96). Detailed protocols are described in SI Materials and Methods.

### Yeast microscopy

Detailed protocols for live-cell fluorescence microscopy and transmission electron microscopy preparation and imaging of yeast cells are described in SI Materials and Methods.

### Sucrose gradient sedimentation

Sucrose gradient sedimentation was performed as previously described (15). Detailed protocols are described in SI Materials and Methods.

## Data Availability

All data is presented within this article and supplementary information.

## Acknowledgements

This work was supported by an NIH grant to DJG (R01GM124216) and an NIH Postdoctoral Fellowship to SLB (F32GM139247). This study was also supported by the Robert P. Apkarian Integrated Electron Microscopy Core (RPAIEMC), which is subsidized by the Emory University School of Medicine and the Emory College of Arts and Sciences. Additional support was provided by the Georgia Clinical & Translational Science Alliance of the National Institute of Health under award number UL1TR000454. Some of the data reported here were collected on the JEOL JEM1400 TEM supported by the National Institutes of Health Grant S10 RR025679. We thank Joan Curcio, Katarzyna Pachulska-Wieczorek, Yury Chernoff, and Pavithra Chandramowlishwaran for providing reagents and advice, and Adam Hannon-Hatfield for valuable discussions and comments on the manuscript.

## SI Materials and Methods

### Plasmids and cloning

Plasmids, primers, and gene fragments are listed in Supplementary Tables 4-6. All Ty1 nucleotide and amino acid information correspond to the Ty1H3 sequence (GenBank M18706.1). All cloning was done with NEBuilder HiFi DNA Assembly Master Mix (New England Biosciences cat. no. E2621). Sup35 fusion plasmids pBDG1691 (1434), pSLBB027 (1134), and pSLBB028 (1258), driven by the *CUP1* promoter, were kindly provided by Y. Chernoff (Chernoff lab plasmid nomenclature in parentheses). Sup35N plasmids contain Sup35 amino acids 1-123, and Sup35NM contains amino acids 1-250. The Gag_PrLD_ contains Gag amino acids 66-136. Gag_PrLD_ fusions to Sup35 were subcloned via EcoRI and XbaI digest and PCR from pBDG598 using primers SLBP0045-7. Hemagglutinin epitope (HA) tags were inserted via XbaI and SacII digest using ssDNA oligos AB42-HA (SLBP0088) or GagPrLD-HA (SLBP0087) and HAtag-SacII (SLBP0086).

pBDG1647 was kindly provided by K. Pachulska-Wieczorek and is the mini-Ty1*his3-*AI plasmid (pJC994) which was constructed by deleting the HpaI-SnaBI fragment of pGTy1*his3AI-[Δ1]* (nucleotides 818-5463 of Ty1-H3) (6).

pBDG1781 contains pGTy1nt.241-5561 which is pEIB (“<u>e</u>nzyme-in-a-<u>b</u>ox”). pEIB was kindly provided by J. Strathern. It was created by deleting the BglII-NcoI fragment which removes the U3 polypurine tract (PPT) and 3’ LTR, preventing reverse transcription of the Ty1 RNA produced from pEIB. The original pEIB provided by J. Strathern also contained a multiply mutated primer binding sequence (PBS), disrupting complementarity to the intitiator tRNA_i_^Met^ which primes reverse transcription. pBDG1781 was corrected back to the original Ty1H3 PBS sequence via XhoI and HpaI digest and PCR from pBDG598 using primers SLBP0116-7.

pBDG1781 derivatives were generated by replacing the Gag_PrLD_ with custom commercial gene fragments (Integrated DNA Technologies (IDT) and Twist Bioscience) via XhoI and HpaI digest. *PrLDΔ* was cloned using SLBG0030 and the chimeras were cloned using overlapping gene fragments SLBG0024, SLBG0025 and a gene fragment encoding the foreign prion domain. The Aβ_1-42_ sequence used is identical to that in pBDG1691 provided by Y. Chernoff and contains a silent mutation at codon 3 (GAA>GAG) to remove an EcoRI site. Mouse PrP (UniProt P04925) amino acid sequence was codon optimized for S. *cerevisiae* using the IDT codon optimization tool.

pBDG1799 contains mature Gag (amino acids 1-401) driven by the *GAL1* promoter fused to GFP-(S65T) with a 7 amino acid linker (nt. CGGATCCCCGGGTTAATTAAC) followed by the *ADH1* terminator sequence, which was kindly provided by J. Curcio on plasmid BJC1066, which is in a pRS415 backbone. The expression construct was subcloned to pRS413 (97) using primers SLBP0221-2 and inserted via XhoI and SacII digest to create pBDG1799. Derivatives were subcloned via XhoI and BbvCI digest and PCR from the corresponding chimeric pEIB plasmids using primers SLBP0117 and SLBP0194.

pBDG598 is pGTy1mhis3-AI, described in (59), and is driven by the *GAL1* promoter and is marked with the *his3*-AI retrotranscript indicator gene. Derivatives were subcloned via XhoI and HpaI digest and PCR from the corresponding chimeric pEIB plasmids using primers SLBP0116-7. All plasmids generated were verified by DNA sequencing.

### SDD-AGE

Semi-denaturing detergent-agarose gel electrophoresis (SDD-AGE) was adapted from published methods (53, 54). Yeast were subcultured from an overnight SC-Ura starter culture into SC-Ura ± 100 μM CuSO_4_ and grown overnight at 30 °C. Approximately 1 x 10^8^ cells were lysed in 200 μL of ice cold Buffer A (50 mM Hepes, pH 7.5; 150 mM NaCl; 2.5 mM EDTA; 1% Triton X-100) with 400 μg/mL PMSF, 16 μg/mL each of aprotinin, leupeptin, pepstatin, and 6 mM DTT by vortexing with glass beads twice for 5 minutes at 4 °C. Cell debris was removed by centrifugation for 2 minutes at 5000 rpm at 4 °C. 4X sample buffer (2X TAE; 20% glycerol; 4% SDS; bromophenol blue) was added to the supernatant and run on a 13 cm 1.8% agarose gel containing 1X TAE and 0.1% SDS at 50 V for several hours until the dye front reached 1 cm from the bottom of the gel. Proteins were transferred to PVDF using 1X TBS by downward capillary transfer overnight at room temperature. The membrane was immunoblotted by standard immunoblotting.

### Ty1*his3-AI* mobility assays

Ty1 retromobility events were detected using the *his3-AI* retromobility indicator gene (59) by qualitative and quantitative assays (58). Qualitative assays were printed from glucose plates onto galactose plates, grown for 48 h at 22 °C, then printed to glucose plates lacking histidine and grown at 30 °C. Quantitative retromobility frequencies were determined from galactose inductions diluted in water, plated on synthetic dropout media, and colonies counted. All experiments were galactose-induced for 48 h at 22 °C. Data represent at least 8 independent galactose inductions; *p*-values were calculated by two-sided Student’s *t*-test. Determination of likely cDNA recombinants versus likely genomic insertions was conducted on His^+^ papillae isolated after 48 hr galactose induction. The *URA3*-bearing plasmid was counter-selected by growth on media containing 5-fluoroorotic acid. Cells that had lost the *TRP1*-bearing plasmid after single colony passaging on YPD were determined by printing to SC-Trp plates. Ura^-^/Trp^-^ cells were tested for growth on SC-His. Loss of the His^+^ phenotype concomitant with plasmid loss indicates a likely cDNA recombinant since the only Ty1 sequence present for homologous recombination is on the plasmids. Retention of the His^+^ phenotype indicates a likely genomic insertion. *p*-values were calculated by Fisher’s exact test compared to wildtype. 100 retromobility events was selected for feasibility of data collection after estimating required sample size of 126 by *a priori* power analysis to detect increased recombination frequency of a 10% effect size with 80% power compared with a 5% recombinant frequency in wildtype piloted with 20 retromobility events. Power analysis for Fisher’s exact test was performed using G*Power 3.1 (98).

### Immunoblotting

Total yeast protein was prepared by trichloroacetic acid (TCA) precipitation using standard techniques (58, 96). Briefly, cells were broken by vortexing in the presence of glass beads in 20% TCA and washed in 5% TCA. Proteins were separated on 8% or 10% SDS-PAGE gels. PVDF membranes were immunoblotted with antibodies at the following dilutions in 2.5% milk-TBST: mouse monoclonal anti-HA antibody clone 2-2.2.14 (Invitrogen cat. no. 26183) (1:1000), mouse monoclonal anti-TY tag antibody clone BB2 (kindly provided by S. Hajduk) (1:10,000) (99), mouse monoclonal anti-IN clone 8B11 (kindly provided by J. Boeke) (1:1,000), rabbit polyclonal anti-RT (Boster Bio cat. no. DZ33991) (1:500), or mouse monoclonal anti-Pgk1 antibody clone 22C5D8 (Invitrogen cat. no. 459250) (1:1000). Immune complexes were detected with WesternBright enhanced chemiluminescence (ECL) detection reagent (Advansta cat. no. K-12049-D50). All imaging was done using a ChemiDoc MP (Bio-Rad). Precision Plus Kaleidoscope protein standards (Bio-Rad cat. no. 1610395) were used to estimate molecular weights.

### Live cell fluorescence microscopy

Following 24 or 48 hr galactose induction, cells were imaged directly in growth media on positively charged slides (Globe Scientific cat. no. 1358W) using a Zeiss Axio Observer.Z1 epifluorescence microscope equipped with an AxioCam HSm camera and captured using AxioVision v4.8.2 software (Carl Zeiss Microscopy).

### Sucrose gradient sedimentation

Following 48 hr galactose induction, a 100 mL culture was harvested, and cells were broken in 15 mM KCl, 10 mM HEPES-KOH, pH 7, 5 mM EDTA containing RNase inhibitor (100 U/mL), and protease inhibitors (16 μg/mL aprotinin, leupeptin, pepstatin A and 2 mM PMSF) in the presence of glass beads. Cell debris was removed by centrifuging the broken cells at 10,000 x g for 10 min at 4°C. Clarified whole cell extract in 500 μL of buffer was applied to a 7-47% continuous sucrose gradient and centrifuged using an SW41 Ti rotor at 25,000 rpm (77,000 x g) for 3 hr at 4°C. After centrifugation, 9 x 1.2 mL fractions were collected, and input and fractions were immunoblotted with TY-tag antibody to detect Gag. Densitometric analysis was performed using Image Lab (Bio-Rad, v. 6.0.1).

### Transmission electron microscopy preparation and imaging of yeast cells

Following 48 hr galactose induction, or 24 hr induction for GFP-expressing strains, cells were fixed with 4% formaldehyde -2.5% glutaraldehyde in 0.1 M sodium cacodylate pH 7.4 for 2 hr at 4 °C, washed three times with cold PBS, once with cold 0.1 M KPO_4_ (pH 6.5), and once with cold P solution (1.2 M sorbitol, 0.1 M KPO_4_ pH 6.5). Cells were spheroplasted in P solution with 25 mM DTT for 15 min at 37 °C using 400 μg/mL of Zymolyase-20T. Spheroplasts were gently washed three times with cold PBS, stored in 0.1 M sodium cacodylate pH 7.4 at 4 °C, and transported to the Robert P. Apkarian Integrated Electron Microscopy Core. Then the cells were washed in fresh 0.1 M cacodylate buffer and spun for 5 minutes at 8000 rpm on an Eppendorf Centrifuge 5430. The cells were spun between each step and were processed in the microcentrifuge tubes in which they were received. After two 0.1 M cacodylate buffer washes of ten minutes each, the cells were post-fixed for an hour in 1% buffered osmium tetroxide. Following two ten-minute washes in distilled water, the cells were en-bloc stained with 0.5% Uranyl Acetate in 0.1 M sodium acetate for 30 minutes. The cells were washed in distilled water for 10 minutes, and then dehydrated in an ascending ethanol series of 15-minute steps starting with 25% and ending with 100% ethanol followed by two 15-minute steps of propylene oxide (PPO). The cells were infiltrated with Eponate12 (Ted Pella, Inc.) epoxy resin in four steps: 1:2 of resin to PPO, 1:1 resin to PPO, 2:1 resin to PPO, and two changes of 100% resin. All resin steps were for 4 hours to overnight, followed by a final change of fresh 100% resin. The cells in resin were then polymerized for two to three days at 60 °C. After release from the tubes, the sample blocks were faced. Ultrathin sections of 70 to 80 nm were made using a Reichert Ultracut S and a Diatome diamond knife. The sections were collected onto 200 mesh copper grids with Carbon stabilized Formvar™ support film then post-stained with 5% Uranyl Acetate and Reynold’s Lead Citrate. Images were acquired using an Ultrascan 1000, 2K x 2K CCD digital camera, on a JEOL JEM1400 TEM operated at 80kV. Micrographs of 140-500 cells per strain were analyzed and representative images were selected for publication. Particle diameters were measured single-blind using FIJI version 2.3.0 (100) by counting at least 60 particles from all cells visible in the field of view (1-3 cells) in at least two separate micrographs.

---

**Fig S1.**
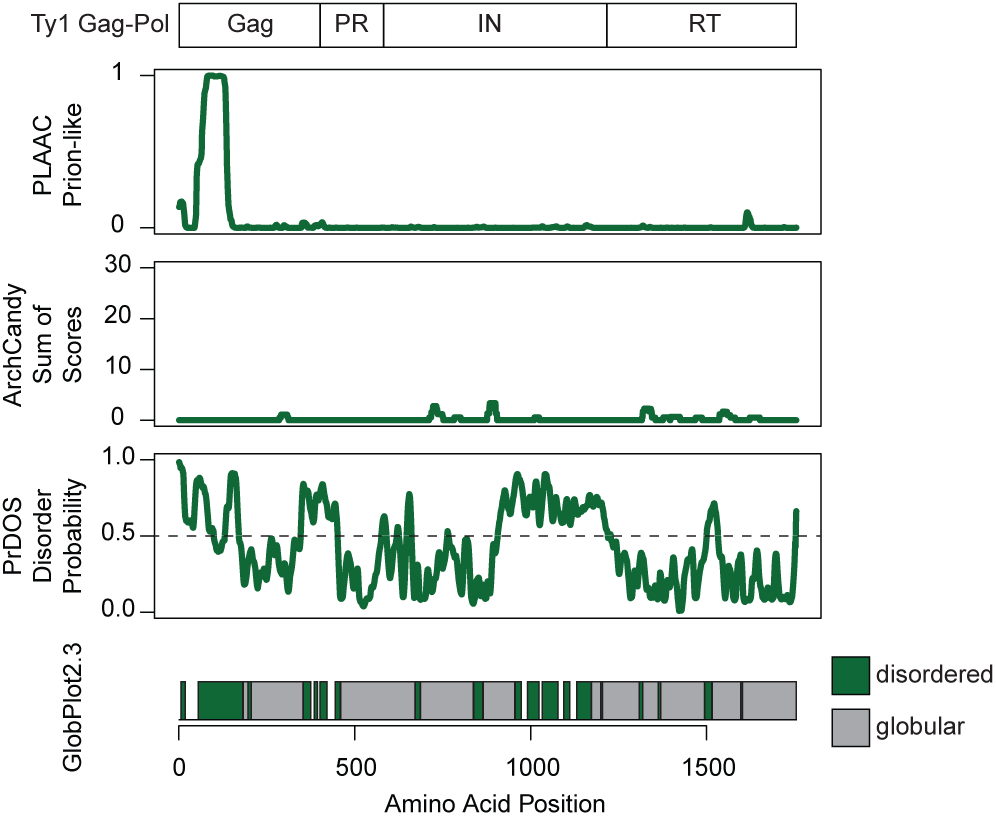
PrLD predictions for Ty1. Schematic of the Ty1 Gag-Pol p199 polyprotein (*top*). Below are bioinformatic analyses aligned with the schematic above: yeast prion-like amino acid composition (PLAAC), predicted amyloidogenic regions (ArchCandy), predicted protein disorder (PrDOS), predicted disordered (green) and globular (grey) regions (GlobPlot2.3).

**Fig S2.**
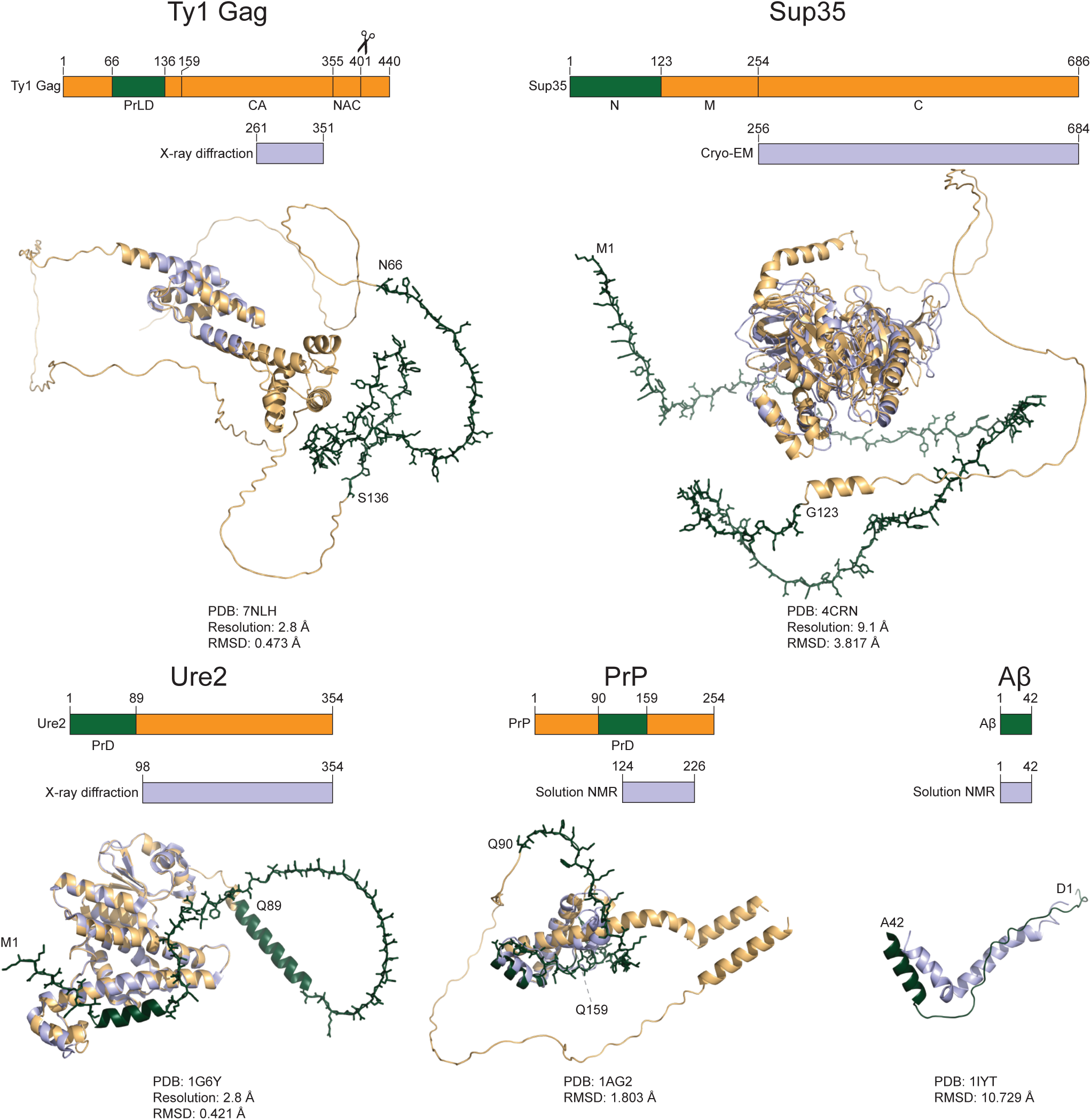
Prionogenic domains are intrinsically disordered in experimental and predicted protein structures. Schematics of protein domains (*top*) and experimentally determined protein structures with the methodology are noted (*bottom*). Amino acid coordinates are shown above cartoon representations of structures predicted by AlphaFold (orange) aligned to published structures (blue). Prion domains are colored in dark green, and their predicted disordered loops are shown in stick representation to aid visualization. PDB accession numbers and reported resolutions for published structures, and RMSD over the common residues between the published and predicted structures, are indicated.

**Fig S3.**
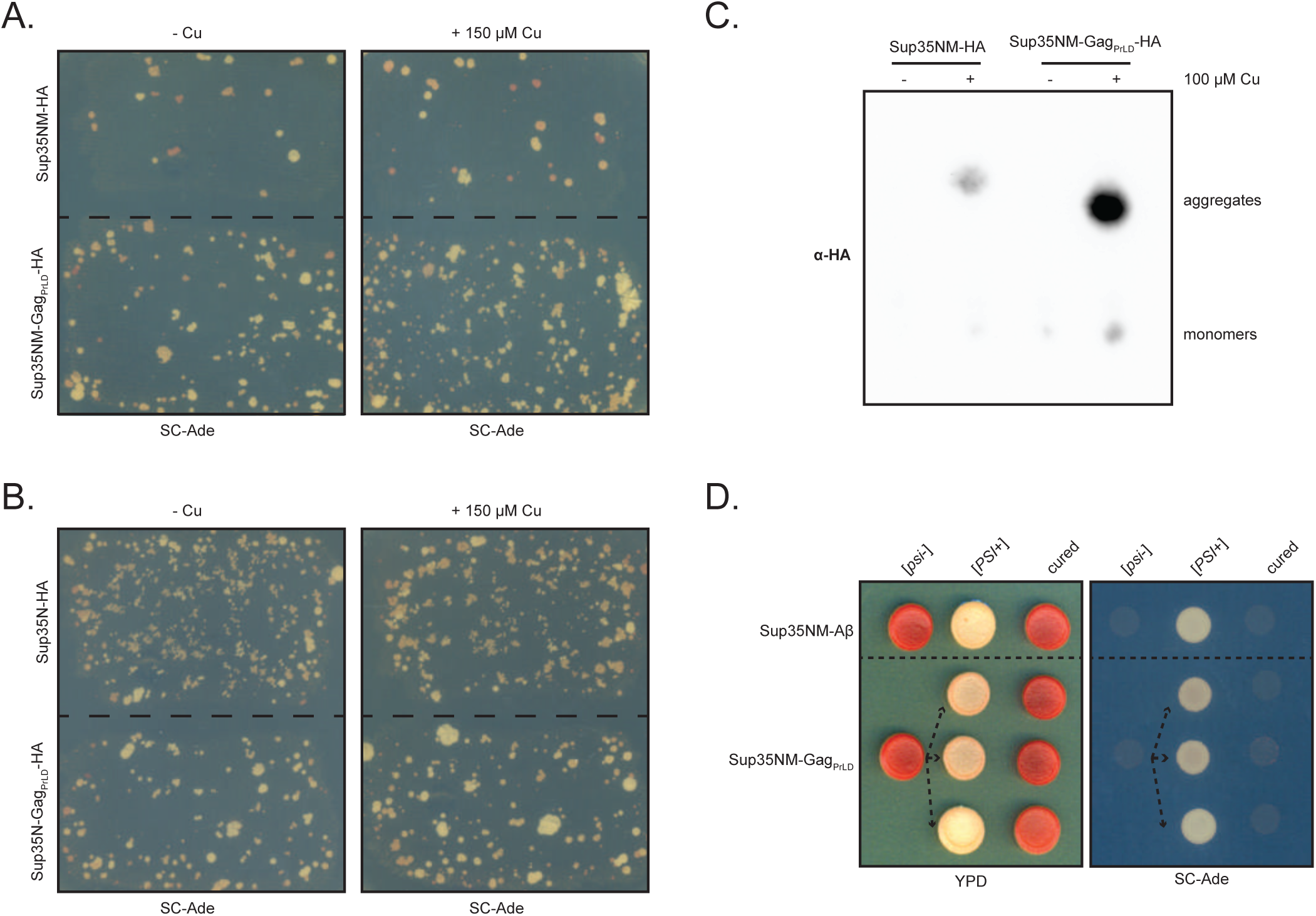
Gag_PrLD_ nucleates a Sup35-based prion reporter. (*A and B*) Qualitative prionogenesis of Sup35 fusions; growth on SC-Ade indicates either a suppressor mutation or [*PSI^+^*] prionogenesis. Expression of Sup35 fusions were induced with 150 μM CuSO_4_. A representative image of at least 3 experiments is shown. (*C*) SDD-AGE analysis of Sup35NM-HA with and without Gag_PrLD_ fusion. Expression of Sup35 fusions were induced with 100 μM CuSO_4_. Monomers and high-molecular weight aggregates of chimeric proteins were detected with anti-HA antibody. A representative image of at least 3 experiments is shown. (*D*) Curing of Ade+ colonies by guanidine hydrochloride (GdHCl) of Sup35NM chimeras. One [*psi^-^*] Sup35NM-Aβ fusion control strain is shown induced to [*PSI^+^*] and cured. Three independent inductions of a [*psi^-^*] Sup35NM-GagPrLD fusion are shown induced to [*PSI^+^*] and cured. [*PSI^+^*] yeast cells are white on YPD and grow on SC-Ade while [*psi^-^*] and cured cells are red on YPD and do not grow on SC-Ade.

**Fig S4.**
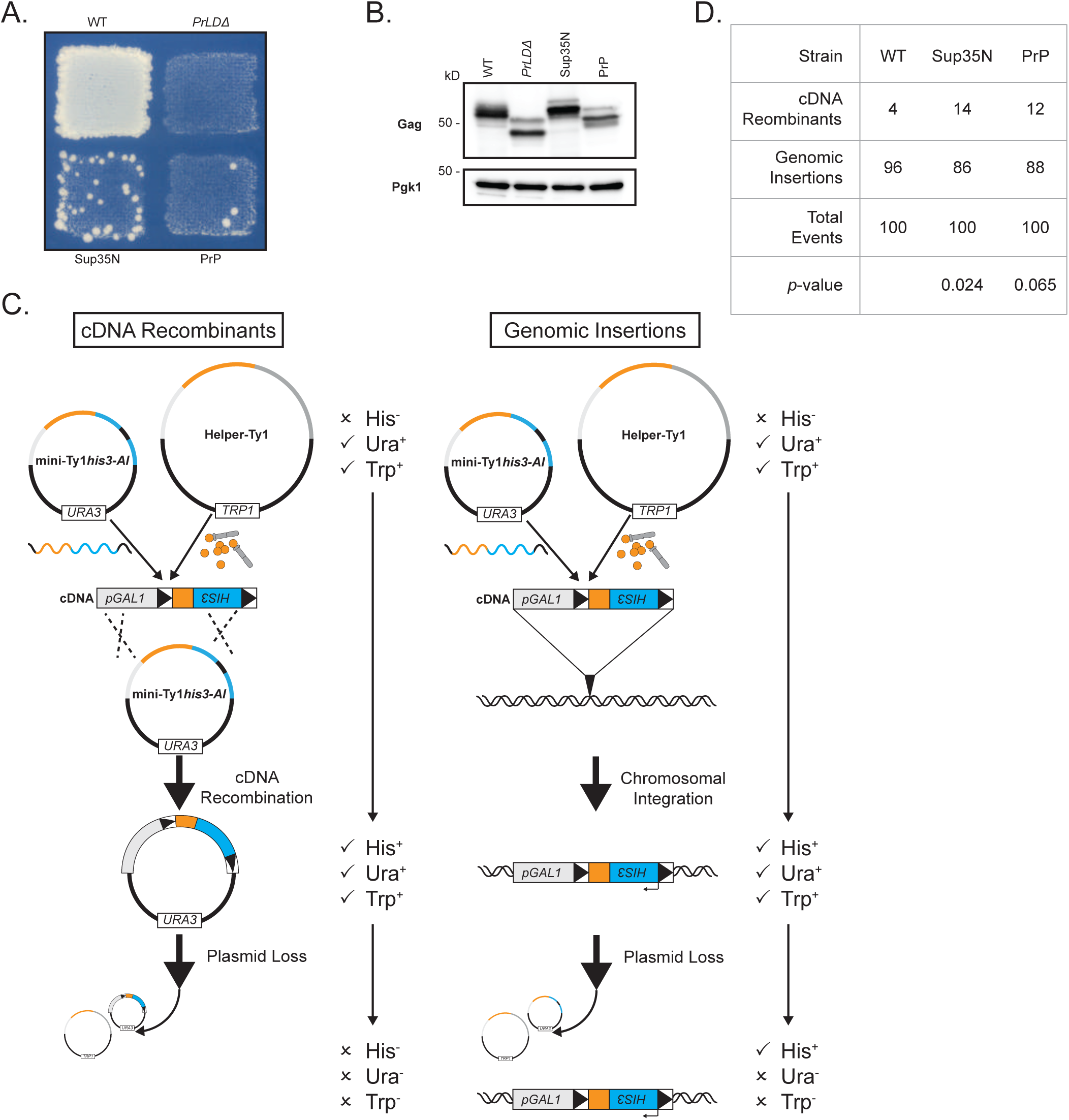
Gag chimeras likely disrupt Ty1 RNA functions and modestly increase cDNA recombination with plasmid-borne mini-Ty1*his3-AI*. (*A*) Qualitative retromobility of chimeric Gag constructs in a single pGTy1*his3-AI* plasmid. Growth on media lacking histidine indicates a retromobility event. A representative image of at least 3 replicates is shown. (*B*) Protein extracts prepared from galactose-induced yeast cells expressing the indicated Gag constructs in a single plasmid were immunoblotted for Gag. Pgk1 serves as a loading control. Migration of molecular weight standards is shown alongside the immunoblots. A representative image of at least 3 replicates is shown. (*C*) Schematic of two major retromobility pathways that lead to His^+^ cells detected in retromobility assays using Ty-less strains. cDNA recombination and genomic insertion can be differentiated by allowing for plasmid loss after a retromobility event and testing for the retention of growth on medium lacking histidine. (*D*) Table indicating the ratio of cDNA recombinants versus genomic insertions, *p*-values are compared to wildtype.

**Fig S5.**
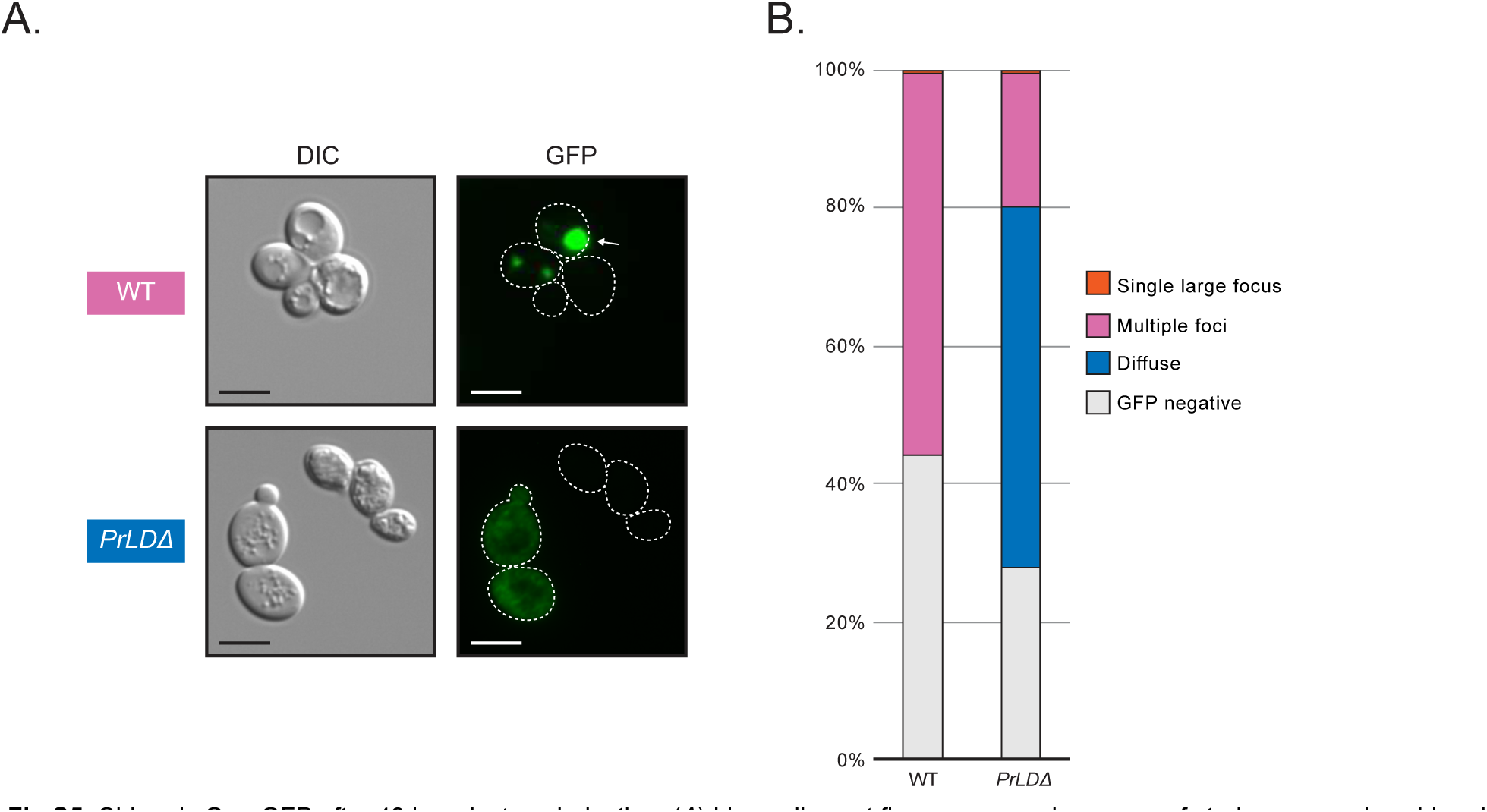
Chimeric Gag-GFP after 48 hr galactose induction. (*A*) Live-cell yeast fluorescence microscopy of strains expressing chimeric Gag-GFP after 48 hr galactose induction. Normaski (DIC) and GFP channels are shown with cell outlines added to GFP channels based on DIC images. The strain labels are colored to match the most common foci observed. White arrows indicate cells with a single large focus. Scale bars represent 5 μm. (*B*) Quantitation of categories of foci observed as a percentage. The multiple foci category includes cells with multiple large foci, one or more small foci, or a combination of both sizes. Exact cell counts are provided in Supplementary Table 2.

**Fig S6.**
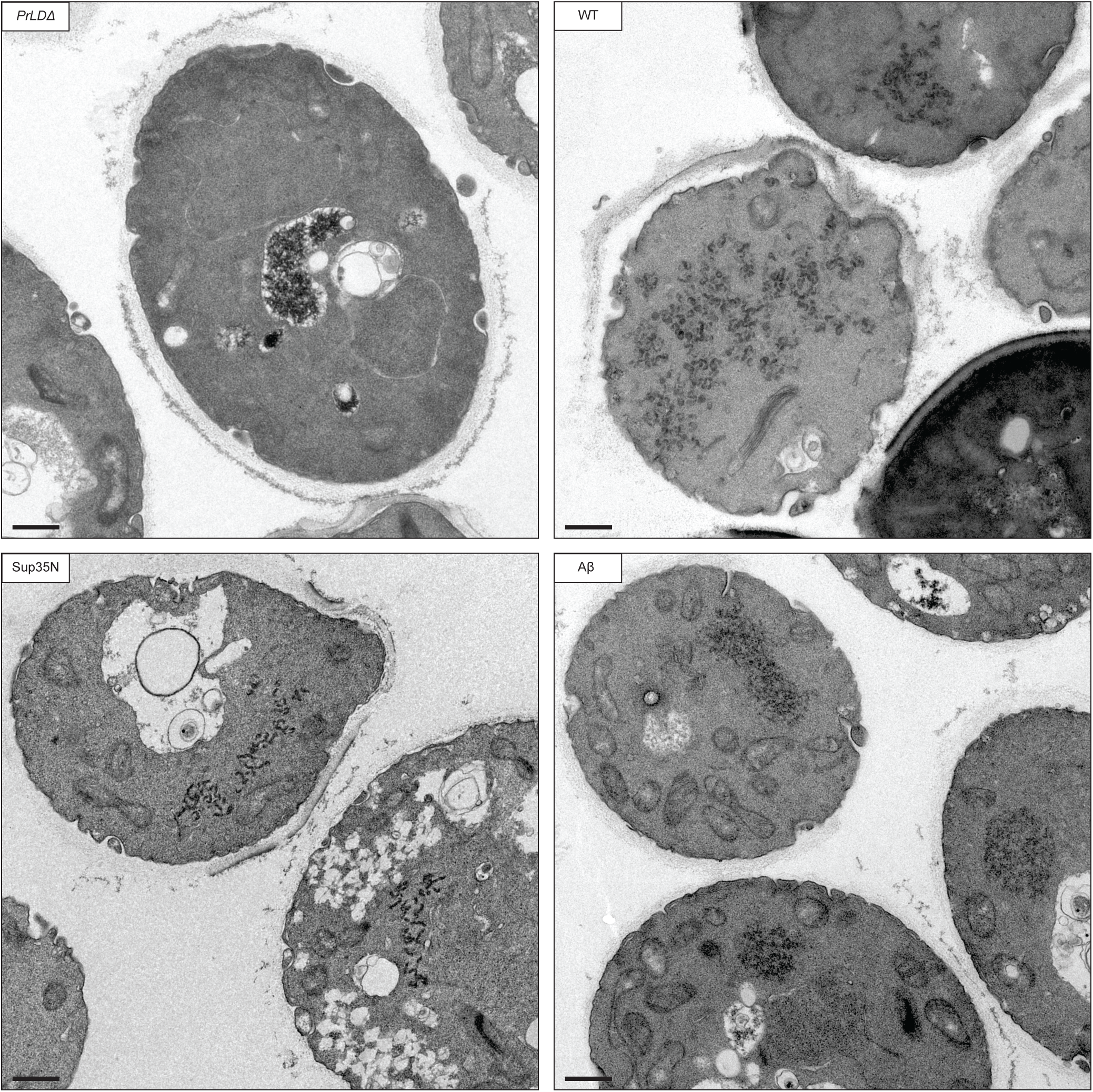
Thin-section TEM of Gag-GFP strains. Thin-section TEM of 24 hr galactose-induced cells expressing Gag-GFP chimeras. Representative cells are shown. Scale bars represent 500 nm.

**Fig S7.**
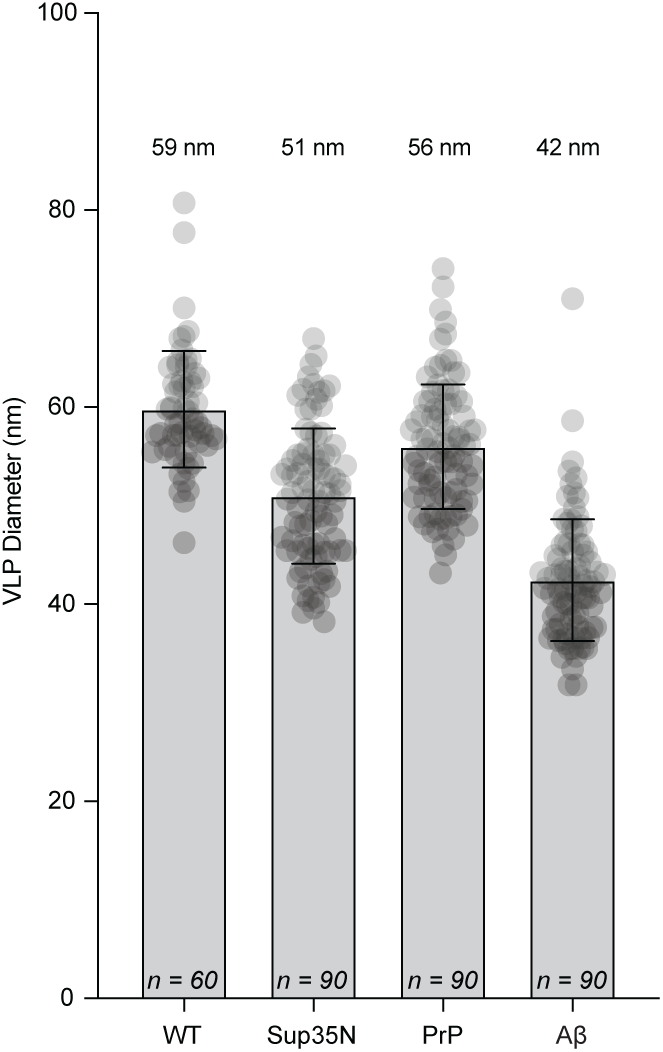
VLP diameter of Gag-PrLD chimeras. Diameter measurements of particles in galactose-induced cells expressing Gag chimeras visualized by thin-section TEM. Each bar represents the mean diameter, displayed as points, and the error bar ± the standard deviation. The median diameter is noted above each bar, the number of particles measured is noted at the base of each bar. Particles from all strains are significantly smaller as calculated from a two-sided Student’s *t*-test compared with WT.

**Supplementary Table S1.** Retromobility frequencies.

| Strain | Label | Retromobility<br>Frequency | Std Dev | *p*-value<sup>a</sup> | Biological<br>replicates |
| --- | --- | --- | --- | --- | --- |
| DG4457 | WT | $6.46 \times 10^{-5}$ | $2.72 \times 10^{-5}$ | Reference | 20 |
| DG4197 | *PrLDΔ* | $8.75 \times 10^{-9}$ | $2.47 \times 10^{-8}$ | $4.75 \times 10^{-7}$ | 8 |
| DG4198 | Sup35N | $6.57 \times 10^{-5}$ | $3.59 \times 10^{-5}$ | 0.933 | 8 |
| DG4201 | Ure2 | 0 | 0 | $4.74 \times 10^{-7}$ | 8 |
| DG4242 | PrP | $1.22 \times 10^{-6}$ | $5.59 \times 10^{-7}$ | $6.51 \times 10^{-7}$ | 8 |
| DG4241 | Aβ | 0 | 0 | $4.74 \times 10^{-7}$ | 8 |

<sup>a</sup> Calculated by two-sided Student's *t*-test

**Supplementary Table S2.** Gag-GFP chimera fluorescent microscopy cell counts.

| Strain | Label | GFP<br>negative | Diffuse | Multiple<br>foci<sup>a</sup> | Single large<br>focus | Total<br>cells |
| --- | --- | --- | --- | --- | --- | --- |
| *24 hr induction* |  |  |  |  |  |  |
| DG4513 | WT | 32 | 0 | 307 | 2 | 341 |
| DG4514 | *PrLDΔ* | 48 | 293 | 50 | 0 | 391 |
| DG4515 | Sup35N | 24 | 4 | 276 | 5 | 309 |
| DG4516 | Ure2 | 38 | 320 | 7 | 0 | 365 |
| DG4517 | PrP | 44 | 45 | 196 | 48 | 333 |
| DG4518 | Aβ | 65 | 23 | 127 | 89 | 304 |
| *48 hr induction* |  |  |  |  |  |  |
| DG4513 | WT | 108 | 0 | 135 | 1 | 244 |
| DG4514 | *PrLDΔ* | 83 | 159 | 58 | 1 | 301 |

<sup>a</sup> This category includes multiple large foci, one or more small foci, or a combination of both sizes.

**Supplementary Table S3.**
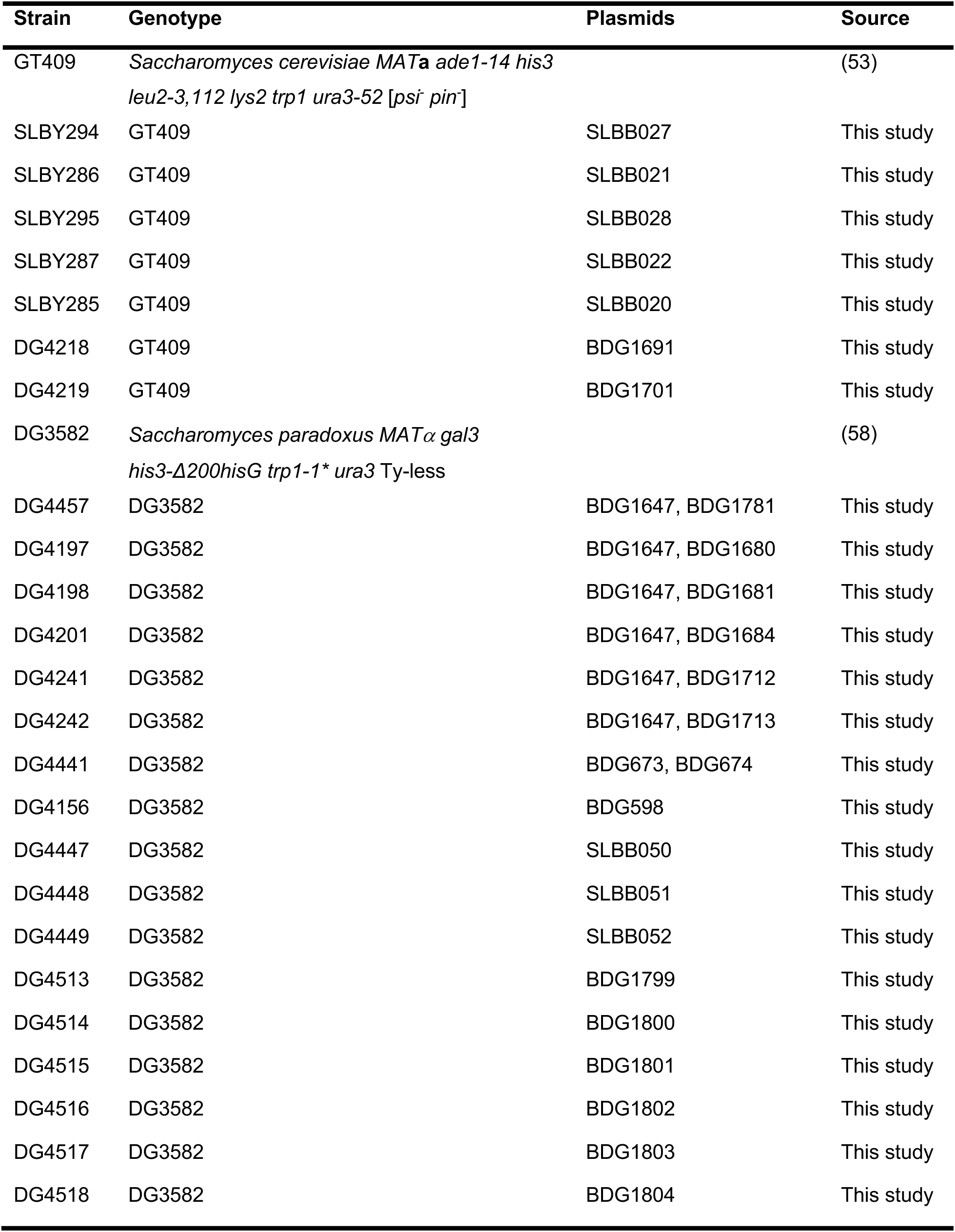
Yeast strains used in this study.

**Supplementary Table S4.** Plasmids used in this study.

| Plasmid | Description | Markers | Source |
| --- | --- | --- | --- |
| pBDG598 | pGTy1*his3-AI* | *URA3/2μ* | (59) |
| pSLBB050 | pBDG598-*PrLD*Δ | *URA3/2μ* | This study |
| pSLBB051 | pBDG598-Sup35N<sub>2-123</sub> | *URA3/2μ* | This study |
| pSLBB052 | pBDG598-PrP<sub>90-159</sub> | *URA3/2μ* | This study |
| pBDG1647 | pGTy1*hisAI*-Δnt818-5463 | *URA3/2μ* | (6) |
| pBDG1781 | pGTy1nt.241-5561 | *TRP1/2μ* | This study |
| pBDG1680 | pBDG1781-*PrLD*Δ | *TRP1/2μ* | This study |
| pBDG1681 | pBDG1781-Sup35N<sub>2-123</sub> | *TRP1/2μ* | This study |
| pBDG1684 | pBDG1781-Ure2<sub>17-76</sub> | *TRP1/2μ* | This study |
| pBDG1712 | pBDG1781-Aβ<sub>1-42</sub> | *TRP1/2μ* | This study |
| pBDG1713 | pBDG1781-PrP<sub>90-159</sub> | *TRP1/2μ* | This study |
| pBDG673 | pRS424 | *TRP1/2μ* | (97) |
| pBDG674 | pRS426 | *URA3/2μ* | (97) |
| pBDG1691 | pCUP1-SUP35NM-Aβ<sub>1-42</sub> | *URA3/CEN* | (53) |
| pBDG1701 | pCUP1-SUP35NM-Gag<sub>PrLD</sub> | *URA3/CEN* | This study |
| pSLBB020 | pCUP1-SUP35NM-Aβ<sub>1-42</sub>-HA | *URA3/CEN* | This study |
| pSLBB021 | pCUP1-SUP35NM-Gag<sub>PrLD</sub>-HA | *URA3/CEN* | This study |
| pSLBB022 | pCUP1-SUP35N-Gag<sub>PrLD</sub>-HA | *URA3/CEN* | This study |
| pSLBB027 | pCUP1-SUP35NM-HA | *URA3/CEN* | <sup>a</sup> |
| pSLBB028 | pCUP1-SUP35N-HA | *URA3/CEN* | (53) |
| pBDG1799 | pGAL-Gag<sub>1-401</sub>-GFP | *HIS3/CEN* | This study |
| pBDG1800 | pBDG1799-*PrLD*Δ | *HIS3/CEN* | This study |
| pBDG1801 | pBDG1799-Sup35N<sub>2-123</sub> | *HIS3/CEN* | This study |
| pBDG1802 | pBDG1799-Ure2<sub>17-76</sub> | *HIS3/CEN* | This study |
| pBDG1803 | pBDG1799-Aβ<sub>1-42</sub> | *HIS3/CEN* | This study |
| pBDG1804 | pBDG1799-PrP<sub>90-159</sub> | *HIS3/CEN* | This study |

<sup>a</sup> Kindly provided by Y. Chernoff.

**Supplementary Table S5.**
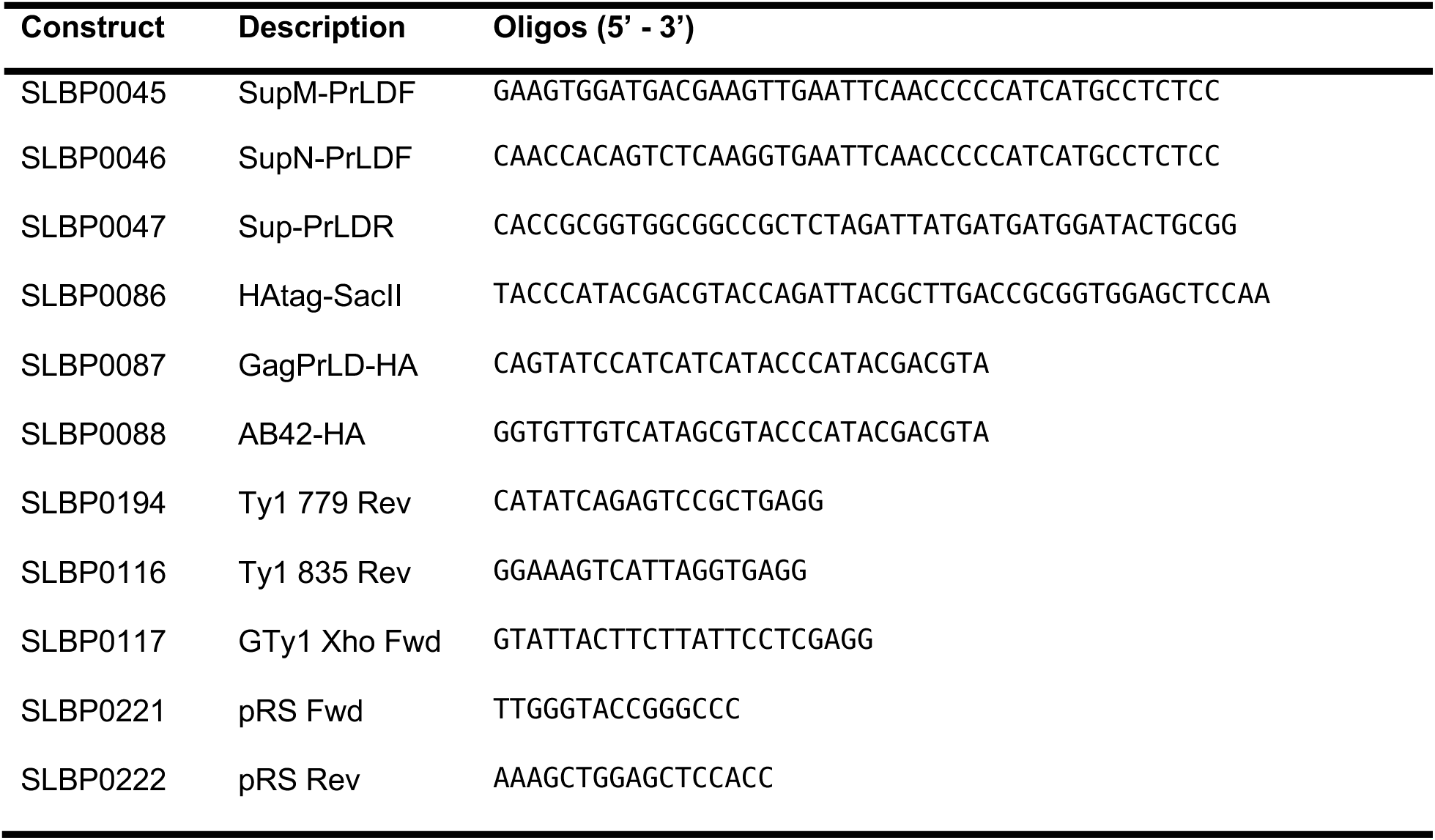
Primers used in this study.

**Supplementary Table S6.**
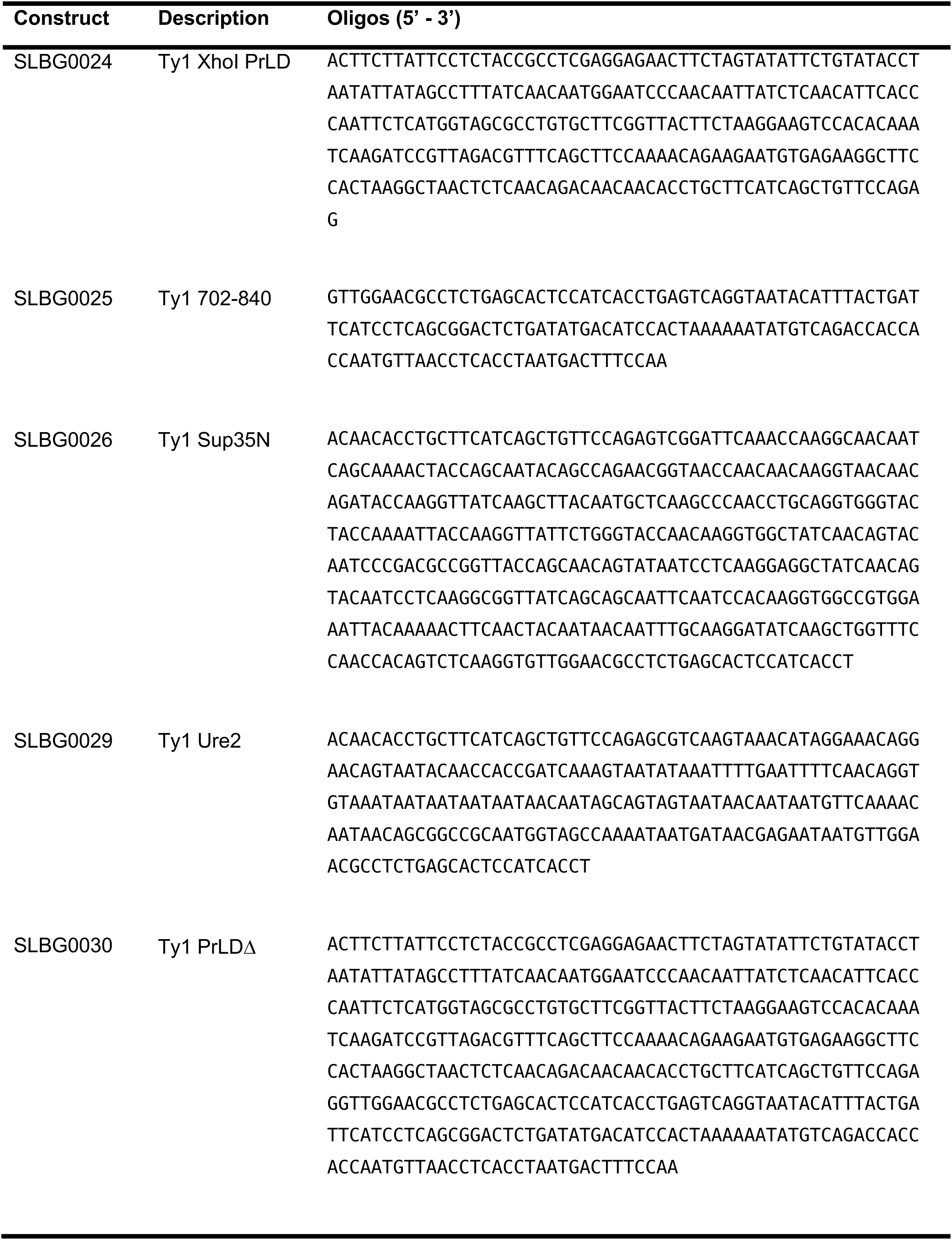
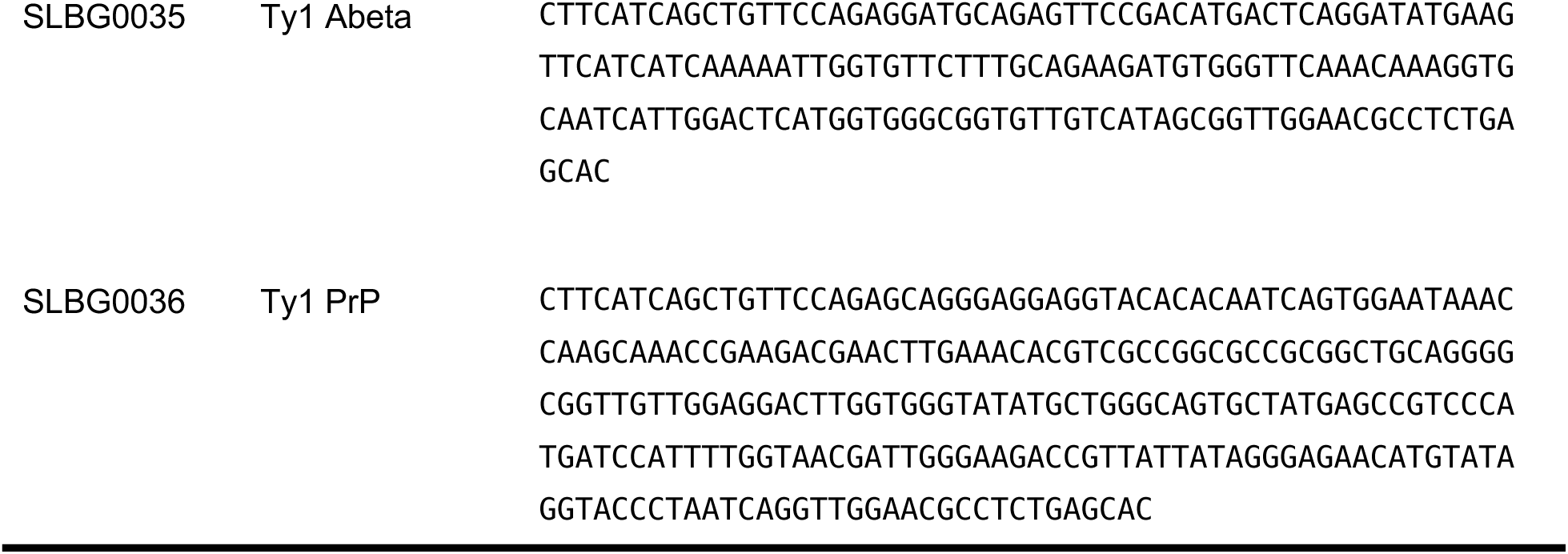
Gene fragments used in this study.

